# Comparative Analysis of TG/CA Repeats in Sixteen Primate Genomes Reveals the Dynamics and Role of TG/CA Repeats in the Human Genome

**DOI:** 10.1101/2023.05.09.539941

**Authors:** Aditya S. Malwe, Samuel Mondal, Pradyumna Harlapur, Vineet K Sharma

## Abstract

Among the different microsatellite sequences found in the human genome, the dinucleotide TG/CA repeats are one of the most abundant, exhibiting multifaceted functional roles. Availability of several primate genomes offers relevant datasets for studying the evolution and function of these repeats in non-human primates and human genome. Using pairwise genomic alignments, genome-wide analysis of these repeats was performed in human and sixteen other primate genomes. The total number of these repeats and expansion of medium (12≤ n< 23) and long (n≥23) (TG/CA)_n_ repeats was significantly higher in human than other primates. Further, other dinucleotide repeats like TA were found in the orthologous genomic regions in other primates. Thus, selection, elongation and a selective process of conversion of other dinucleotide repeats in primates to TG/CA repeats in humans was apparent and presented in this study as a comprehensive model for the dynamics and role of TG/CA repeats in the human genome.

## 1. Introduction

Short tandem repeats of DNA consisting of repetitive units 2 to 6 base pairs are commonly referred to as microsatellites. They are widespread in the human genome; and because they exhibit length variability, they are often used as genetic markers in population studies [1, 2]. Several models have been proposed to explain the polymorphic nature of these repeats. One of the earliest models, referred to as the stepwise mutation model (SMM), suggested that mutations change the repeat length by addition or removal of one or more units [3]. Later, a more comprehensive model was proposed which suggested that a genome-wide distribution of microsatellite repeat length that is at equilibrium results from a balance between length and point mutations [4, 5]. It has been proposed that these length changes primarily arise by polymerase slippage during DNA replication [6, 7, 8]. By slippage mutation, shorter repeats expand and longer repeats contract [5, 9]. The minimum number of repeat units needed for expansion has been proposed as six units [9], however some recent reports suggest that there is no such threshold length [10]. The maximum length observed is in the range of 30 repeat units and this limit is rarely exceeded because long repeats are highly unstable and tend to mutate back to shorter lengths. Recombination has also been proposed to play a role in altering the length of microsatellites [11].

Among all the microsatellite repeats of various lengths, dinucleotide repeats are the most abundant. And, among the dinucleotide repeats, (TG/CA)_n_ repeats are the most abundant and most widely distributed in the human genome [4, 8, 12]. These repeats exhibit length polymorphism by DNA replication slippage [13, 14] and also by recombination events [15]. Among all the microsatellites, these repeats have traditionally been exploited to construct genetic maps [1]. However, the most characteristic property of these repeats is their propensity to adopt the unusual Z-DNA conformation under super-helical stress during transcription [16, 17, 18]. Z-DNA act as *cis*-modulators of gene expression and the unusual Z-DNA conformation is stabilised by Z-DNA binding proteins [19,20]. The ability of (TG/CA)_n_ repeats to form Z-DNA conformation enables them to modulate gene expression, and several studies have indicated the role of these repeats as *cis*-modulators of gene expression [16, 21, 22, 23, 24, 25]. A recent study showed that long polymorphic TG repeats can alter methylation of downstream CpG island in goldfish models [26]. (TG/CA) repeats also play a role in intron splicing as shown by a study conducted by Hui et al., where they showed that length of CA repeats can alter splicing efficiency by serving as HnRNP L recruitment site during intron splicing in *eNOS* gene [27]. These repeats have also been shown to act as hot spots of recombination [11]. The abundance and the above-mentioned functional roles of these repeats underscore their significance in the human genome; and thus, their evolution in the human genome becomes an intriguing question.

An earlier study by Cooper et al. showed that out of 19 informative microsatellite markers, consisting mainly of (TG/CA)_n_ repeats, 13 are longer in human as compared to their chimpanzee orthologues. This study also suggested a mutational bias in favor of microsatellite expansions and a higher-than-average genome-wide microsatellite mutation rate in the human lineage [28]. Another study using a small region (5.1 MB) of human and chimpanzee genomic DNA alignments showed that microsatellites are longer in humans as compared to chimpanzees [29]. These studies using limited data show that microsatellites are longer in humans as compared to chimpanzees, the nearest living common ancestor to humans. Thus, by including the genomic sequences from other primates, a comprehensive analysis of (TG/CA)_n_ repeats in primate genomes will serve to illuminate the evolution of these repeats over a wider phylogenetic distance.

The recent availability of genomic sequences of several non-human primates offers relevant datasets for the evolutionary comparison of repetitive sequences. Genome-wide sequence comparison of orthologous locations, where these repeats are conserved in primate genomes due to their common ancestors, allows us to investigate the evolutionary dynamics of these repeats. Here, we present a genome-wide analysis of (TG/CA)_n_ repeats using pair-wise chromosomal alignments of human with 16 other primate genomes. The relative elongation and contraction of these repeats was analysed in the genic and inter-genic regions in the genomic alignments. The role of the repeat flanking regions in the expansion or contraction of these repeats was also examined. From this analysis, we proposed a model to explain the dynamics of the expansion-contraction process of these repeats during the evolution of primates.

## 2. Results

### 2.1. Summary of the genome-wide alignments

The evolutionary tree of the primates analysed in this study is shown in Figure 1. Using the alignments (selected according to the criteria defined in the Methods section), the percentage alignment coverage was found between each of the primate and human genomes. As expected, a higher percentage of alignment in primates closely related to humans was found compared to the primates distantly related to humans (Figure S2), with slight discrepancies observed for most distant primates like Tarsier, Mouse Lemur and Bushbaby, possibly due to their limited genomic studies and less updated assemblies available.

**Figure 1.**
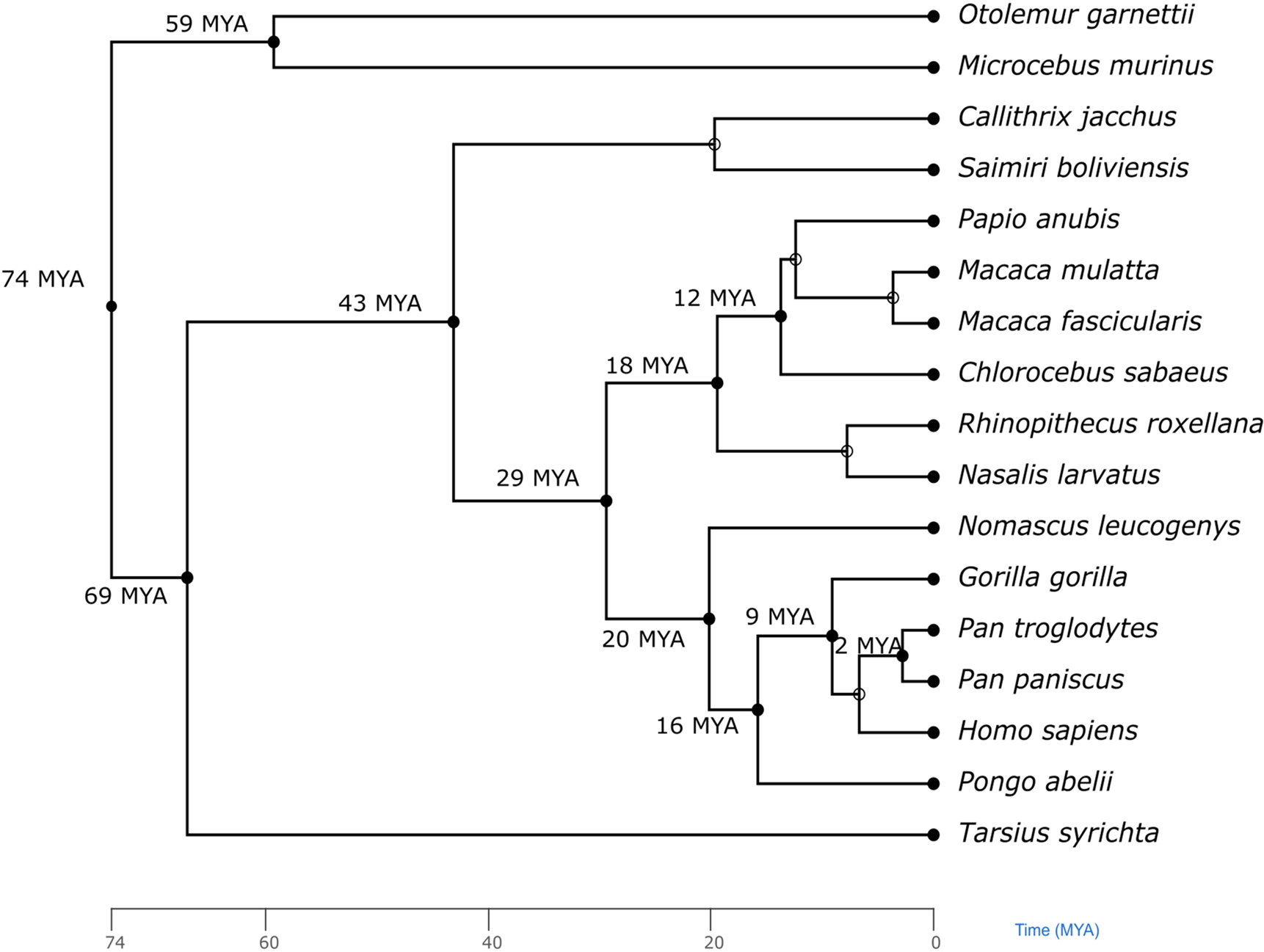
Evolutionary tree of the primate genomes analyzed in this study

### 2.2. (TG/CA)_n_ repeats in genomic alignments

The (TG/CA) _n_ repeats were first mapped in the pairwise genomic alignments and the percentage of orthologous repeats in each of the alignments was calculated. Chimpanzee (82.5%), Bonobo (82.8%) and Gorilla (83.8%) genomes showed the highest percentage of orthologous repeats present in human genome, indicating that the total number of these repeats are higher in humans as compared to the other primates. It was also observed that the percentage of orthologous repeats found in the human and the corresponding primate genome decreased gradually in the order of evolutionary distance in most cases (Supplementary table S1) (Figure 2(a)). The length of the repeats at the orthologous locations of the chromosomal alignments was then examined for elongation, contraction and conservation (referred to as equal-length) of (TG/CA)_n_ repeats in human genome compared to the other primate genomes. A higher proportion (0.51-0.94) of (TG/CA)_n_ repeats were found to be elongated in humans at the same loci as compared to the other primates. Only a small proportion (0.04-0.45) of orthologous repeats were found to be of equal length in the genomes, and this proportion decreased with increasing evolutionary distance. Furthermore, even smaller proportions (0.01-0.05) of these repeats were found to have contracted in the human genome (Figure S1). A direct correlation with increasing evolutionary distance was also observed in elongation of (TG/CA) repeats in human genome when compared to other primate genomes (Figure 2(d), Figure S3) with R = 0.9 (p-value = 0.000001). However, an inverse correlation is seen in case of contraction and conservation of (TG/CA) repeats with R = −0.75 (p-value = 0.0009) and R = −0.89 (p-value = 0.000004) respectively (Figure 2(b), Figure 2(c), Figure S3).

**Figure 2.**
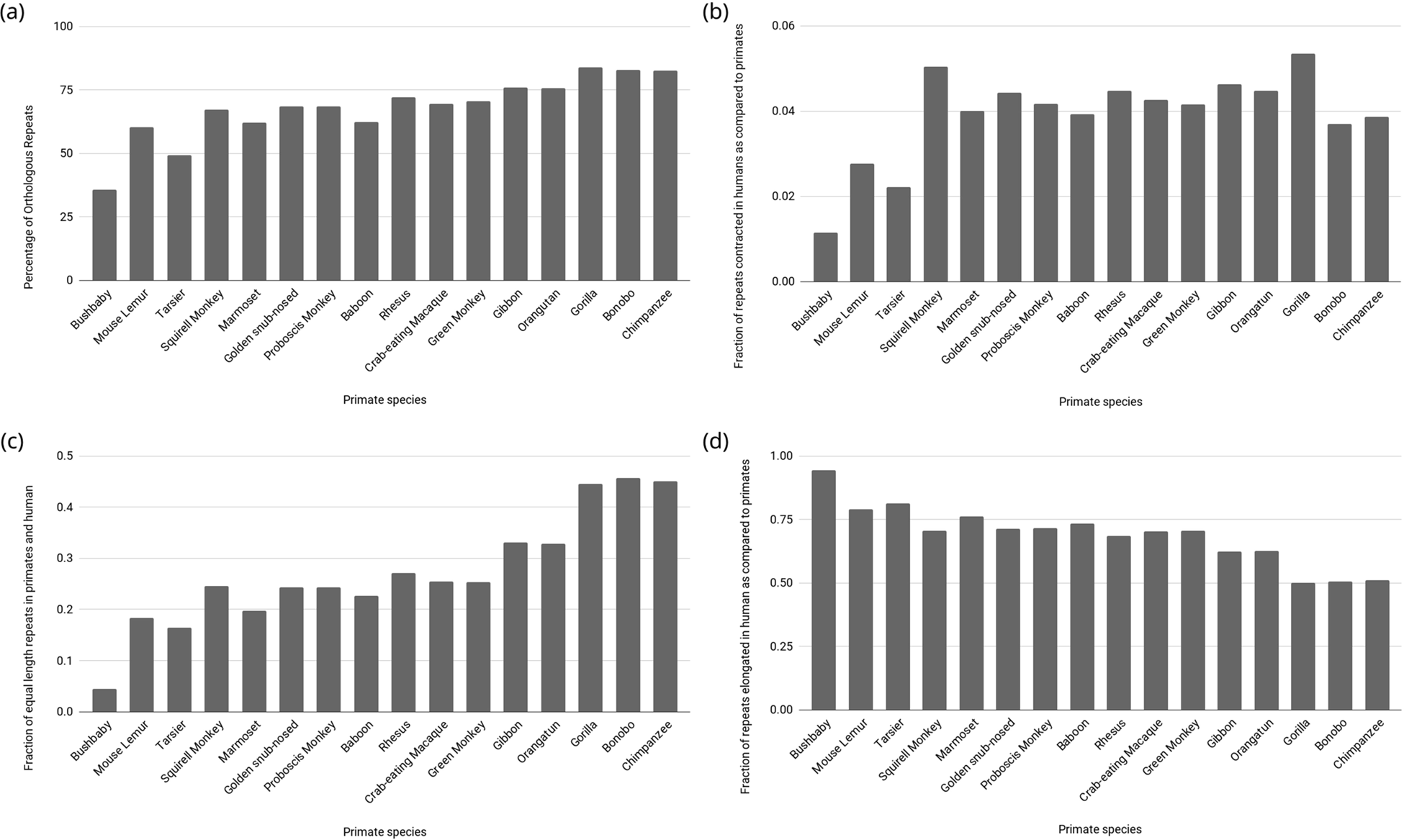
(a) Percentage of (TG/CA)_n_ repeats which were found at orthologous locations in human and other non-human primate genome alignments (b) Fraction of (TG/CA) repeats elongated in humans as compared to primates (R = 0.9, p-value = 0.000001) (c) Fraction of (TG/CA) repeats contracted in humans as compared to primates (R = −0.75, p-value = 0.0009) (d) Fraction of (TG/CA) repeats with equal length in humans and primates (R = −0.89, p-value = 0.000004)

The repeats were classified into three length categories to examine the correlation between repeat length and elongation or contraction frequency: short (Type I), medium (Type II) and long (Type III) (Methods section). Among the repeats which were elongated in humans, a significantly high proportion (0.51-0.63) were Type II except in case of Bushbaby. In addition, higher proportions (0.055-0.198) of Type III repeats were also found in elongated repeats as compared to the proportion of Type III repeats in equal-length (below 0.005) and contracted repeats (below 0.002) (Figure 3). This observation indicates that medium repeats have shown expansions converting to long repeats. Conversely, large proportion of short Type I repeats are observed in case of contracted and equal-length repeats in humans.

**Figure 3.**
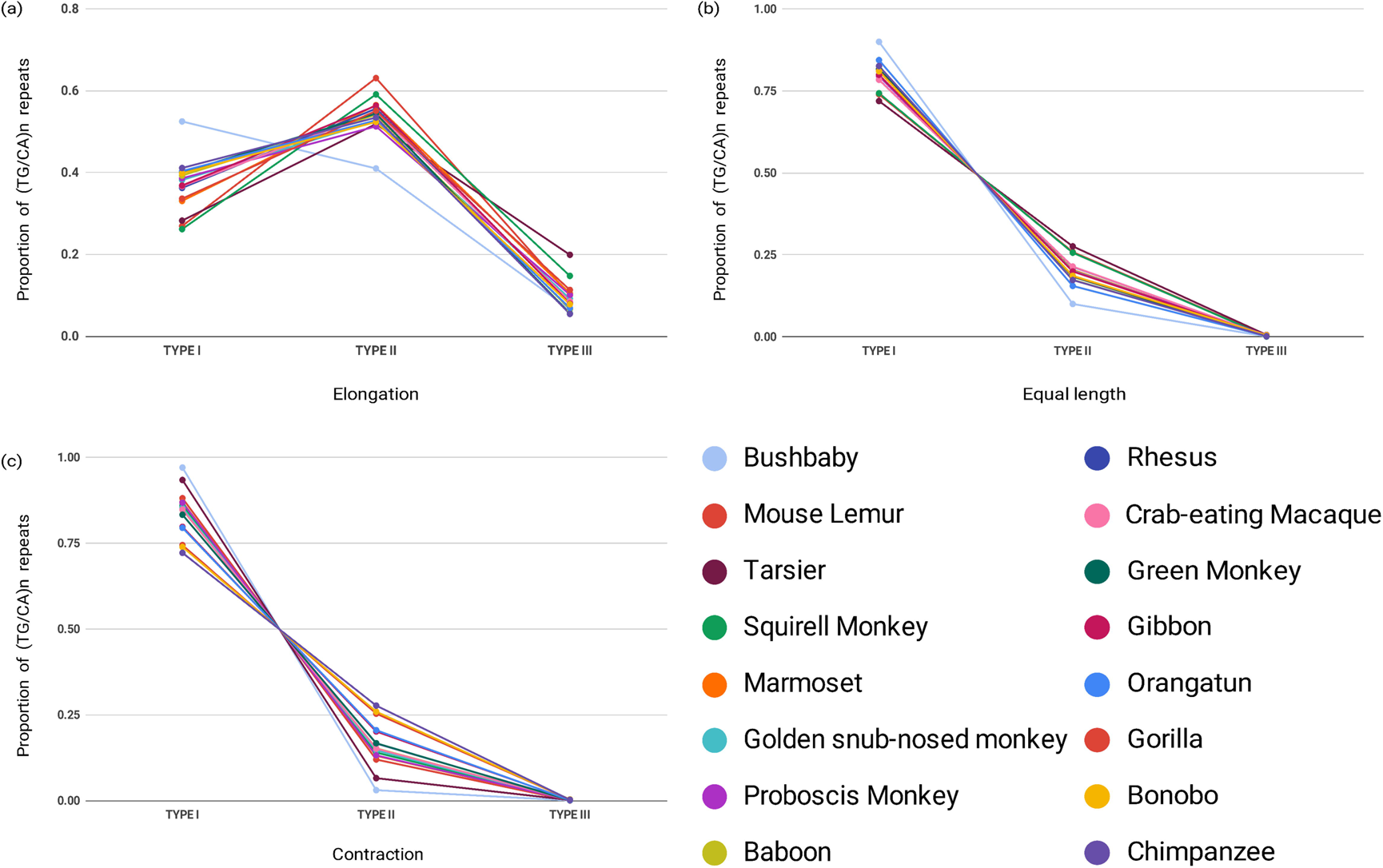
Distribution of the types (I-short, II-medium, III-long) of (TG/CA)_n_ repeats with respect to repeat length (equal-length, elongated, and contracted) in the human genome compared to the other primate genomes.

### 2.3. (TG/CA)n repeats in gene regions

To examine the distribution of (TG/CA)_n_ repeats with respect to gene and intergenic regions, the sequences from the human and chimpanzee alignments were used because chimpanzee is closely related to human, with human and chimpanzee alignments spanning the major part (∼94%) of the human genome. About 75% of (TG/CA)_n_ repeats are present in intergenic region while remaining are present in intragenic regions of human genome. Moreover, around 97% of the intragenic repeats are present in intron and only 3% are present in exon. Proportions of repeats contracted, elongated and equal-length in humans when compared to chimpanzee genome mimics the global distribution. Similarly, proportion of repeats elongated in humans is more across all genomic regions as compared to proportion of repeats contracted in humans. Proportion of repeats with equal length in human and chimpanzee genome is comparable to contracted repeats (Figure 4) (Figure S4).

**Figure 4.**
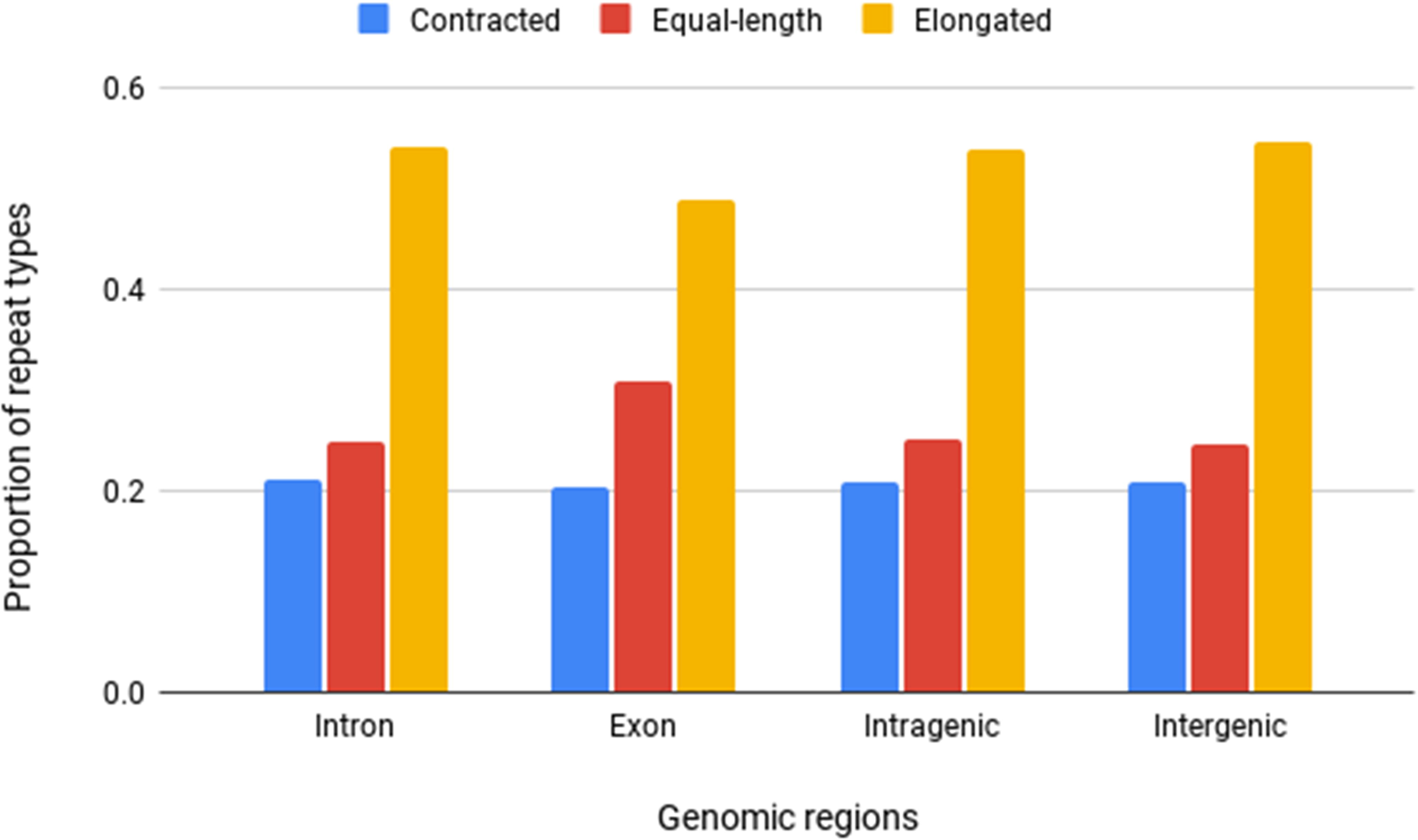
Proportion of contracted, equal-length and elongated repeats in different genomic regions.

### 2.4. Transitions and transversions in the repeat regions

To understand the evolutionary processes underlying the repeat-length dynamics in humans, the frequency of nucleotide transitions and transversions in the (TG/CA)_n_ repeat region was examined. The global genome-wide frequencies of transitions and transversions in the genomic alignments were first calculated. As expected, the genome-wide frequencies of transitions (16.63%) were higher as compared to transversions (4.18%) (Figure S5). In the figure, the direction of change, for example G->A or C->T, is shown from each respective primate to human for a single strand, therefore, both G->A and C->T were considered for the analysis separately instead of combining them together. In all genomic alignments, the frequencies of all four types of transitions (G->A, C->T, A->G and T->C) were very similar (∼15-17%), whereas the frequencies of all types of transversions were less than 6%. The observed genome-wide frequency of transitions and transversions corroborates with their previously reported genome-wide frequencies [8] and suggests that there is no mutational bias for conversion of any nucleotide into another nucleotide.

In contrast to the above, within the alignments spanning the (TG/CA)_n_ repeat regions some interesting trends were observed. In the case of TG repeats, out of the four possible types of transition events only two were observed among which the transition from A->G was the most frequent (35-44%), followed by C->T (20-23%) (Figure 5a). The most frequent transversions were from T->G (12-15%) followed by A->T (6-10%) and C->G (7-10%), while the least common was G->T (5-8%). Similarly, in the case of CA repeats, out of the four possible types of transition events only two were observed among which the transitions from T->C were the most frequent (35-43%), followed by G->A (20-23%) and the most frequent transversions were from A->C (12-15%), T->A (6-10%), G->C (8-10%) and then C->A (5-9%) (Figure 5b). The frequencies of the various transitions and transversions were similar in all genomes for both TG and CA repeat regions. It is evident that the overall frequencies of transitions and transversions in the repeat regions from non-human primate genomes to humans do not follow the genome-wide pattern. Only a few types of transitions and transversions are more frequent as compared to the global frequencies.

**Figure 5.**
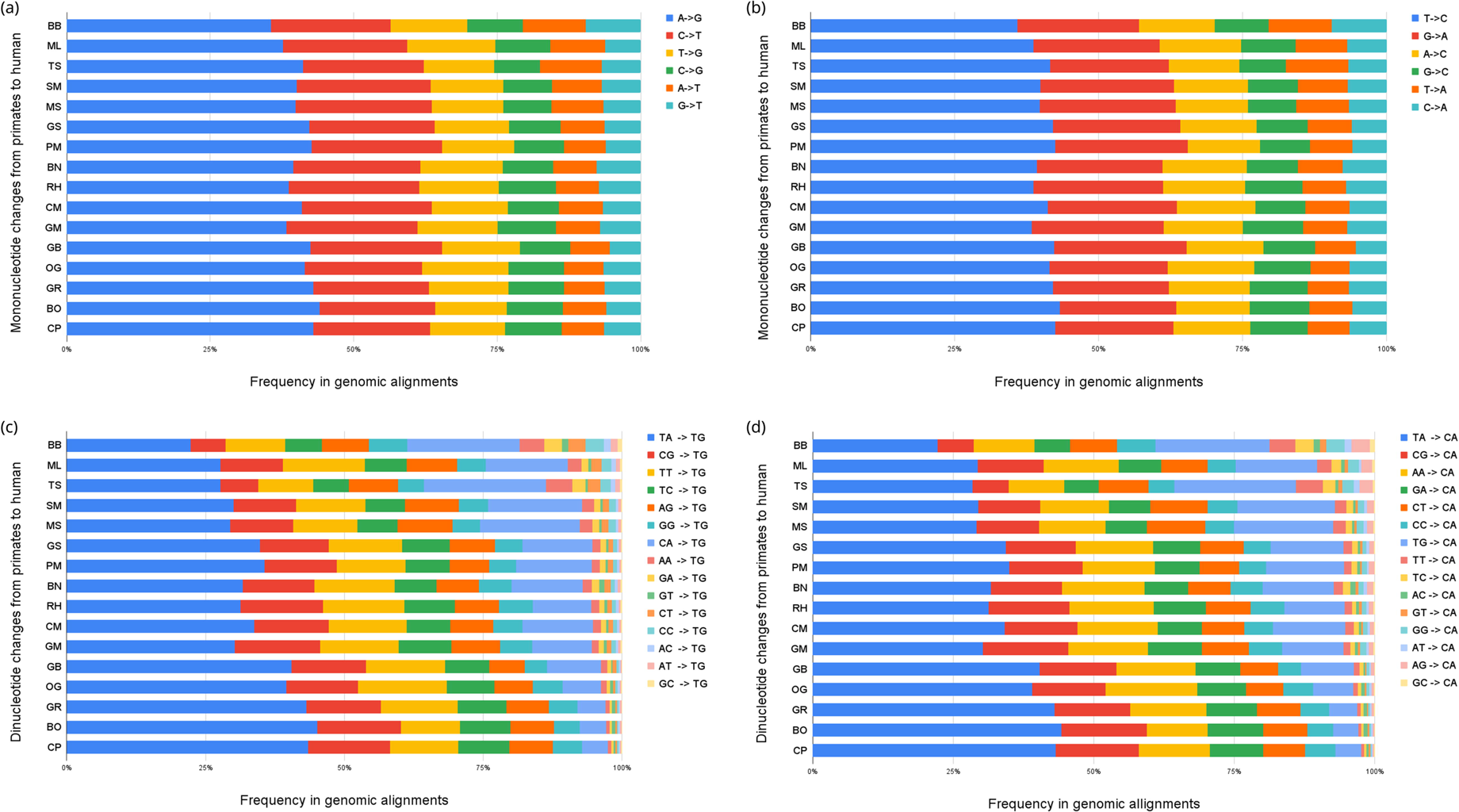
(a) Frequency of transitions and transversions in the TG repeat regions in the human and other primate genomic alignments. (b) Frequency of transitions and transversions in the CA repeat regions in the human and other primate genomic alignments (c) Frequency of transitions and transversions at the dinucleotide level in the TG repeat regions in the human and other primate genomic alignments (d) Frequency of transitions and transversions at the dinucleotide level in the CA repeat regions in human and other primate genomic alignments

To examine the significance of these transitions and transversions at the dinucleotide level in the human and other primate alignments, the dinucleotide conversion frequencies, which in effect result from the above mononucleotide substitutions, were analyzed. Within the TG repeat regions (Figure 5c), TA->TG conversion frequencies were the highest (22-45%), followed by CG->TG (6-15%), TT->TG (9-14%), TC->TG (6-9%) and AG->TG (6-9%), and then GG->TG (4-6%). These frequencies correlate with the above transition and transversion frequencies of the nucleotide being changed. Similarly, within the CA repeat regions (Figure 6d), TA->CA conversion frequencies were the most frequent (22-43%), followed by CG->CA (6-15%), AA->CA (9-16%), GA->CA (6-9%), CT->CA (7-10%) and then CC->CA (4-6%). These observations indicate that the dinucleotide conversion frequencies show the same trend as the mononucleotide frequencies of transitions and transitions in the TG and CA repeat regions. The remaining dinucleotide conversions account for less than 5% changes in total. Together, these observations indicate that in these repeat regions, there is a conversion of other dinucleotides to (TG/CA)_n_ repeats from other primates to humans, which could be one of the reasons for the observed higher number and elongation of these repeats in the human genome.

**Figure 6.**
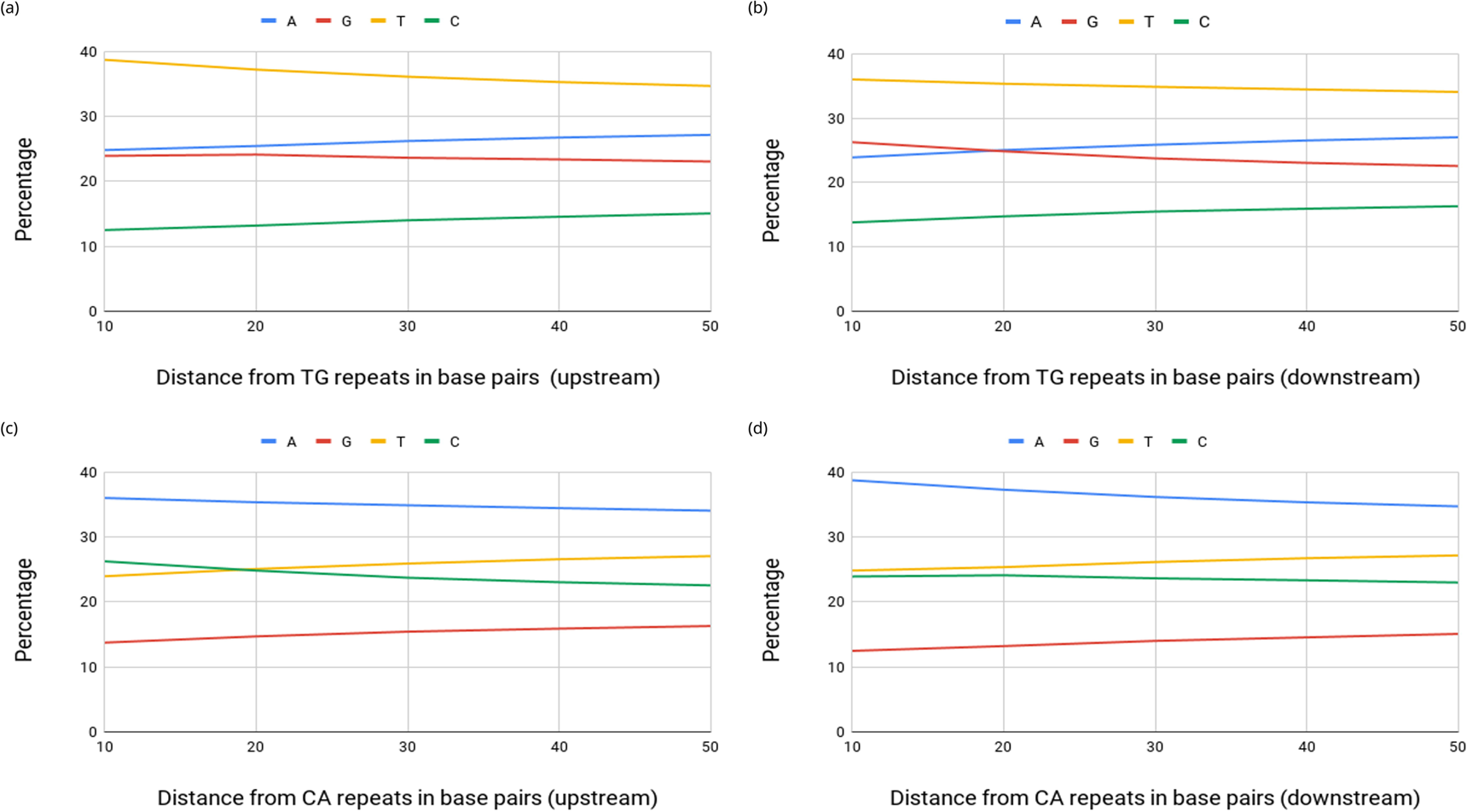
Mononucleotide composition of the 50 bp flanking region upstream and downstream of (TG/CA)_n_ repeats in human.

### 2.5. Size of repeat units undergoing conversion

The analysis to explore the association between the size of repeat unit and its conversion frequency revealed that the conversion of other dinucleotides to TG or CA occurs mostly in single-unit steps; but surprisingly, in several cases long stretches of these dinucleotides were also found to be converted to TG or CA repeats (Supplementary table S2, Supplementary table S3). For both TG and CA repeats in humans, the corresponding orthologous dinucleotide (other than TG/CA) in other primates is most commonly TA, followed by CG and then other dinucleotides. Another interesting pattern that emerged from these tables is that among the 16 primates, the evolutionary distant primates had a dinucleotide repeat consisting of repeat units other than TG or CA at the orthologous location which is now noted as TG or CA at that location. These observations makes it tempting to speculate that there is a directional conversion of other dinucleotide repeats to (TG/CA)_n_ repeats in the human genome during evolution as compared to the other primate genomes.

### 2.6. Composition of the repeat flanking regions

To examine the role of the genomic region flanking the (TG/CA)_n_ repeats on their expansion and contraction, the mononucleotide and dinucleotide composition of the 50 bp flanking region using a 10 bp window was examined in human and chimpanzee genome alignment. It was observed that in the 10 bp region upstream of (TG)_n_ repeats, T was the most frequent (∼36%) base pair followed by G and A and then C (Figure 6a); whereas in the 10 bp region downstream of (TG)_n_ repeats, T was the most abundant (∼39%) but A and G were almost similarly abundant and then C (Figure 6b). However, with increasing distance from 10 to 50 bp, the frequencies of A and C increased, whereas the frequencies of T and G both decreased.

A complementary trend to the above was observed in the case of CA repeats. In the 10 bp region upstream of (CA)_n_ repeats, A was the most frequent (∼36%) base pair but T and C were equally abundant and then G; whereas in the 10 bp downstream region, A was the most frequent (39%) base followed by C and T and then G. However, with increasing distance from 10 to 50 bp, T and G both increased, whereas A and C both decreased (Figure 6c, Figure 6d). Since, TG and CA repeats are reverse complements of one another, the same reverse complementary trend was observed.

At the dinucleotide level, in the regions 10 bp upstream and downstream of the TG (TG)_n_ repeats in the human genome, TG (∼25%) and TA (16-23%) dinucleotides were more abundant. The frequency of these two dinucleotides decreased with increasing distance (10 to 50 bp) from the (TG)_n_ repeats with a sharp decline in case of TG (Figure 7a, Figure 7b). The other dinucleotides, which were less frequent in the 10 bp upstream regions showed an increase in frequency from 10 to 50 bp. Similarly, in the regions 10 bp upstream or downstream of the (CA)_n_ repeats, CA (∼25%) and TA (18-23%) dinucleotides were more abundant (Figure 7c, Figure 7d). The frequency of these two dinucleotides decreased with increasing distance (10 to 50 bp) from the (CA)_n_ repeats with a sharp decline in case of CA. Likewise, the other dinucleotides which were less frequent in the 10 bp upstream regions showed an increase in frequency from 10 to 50 bp.

**Figure 7.**
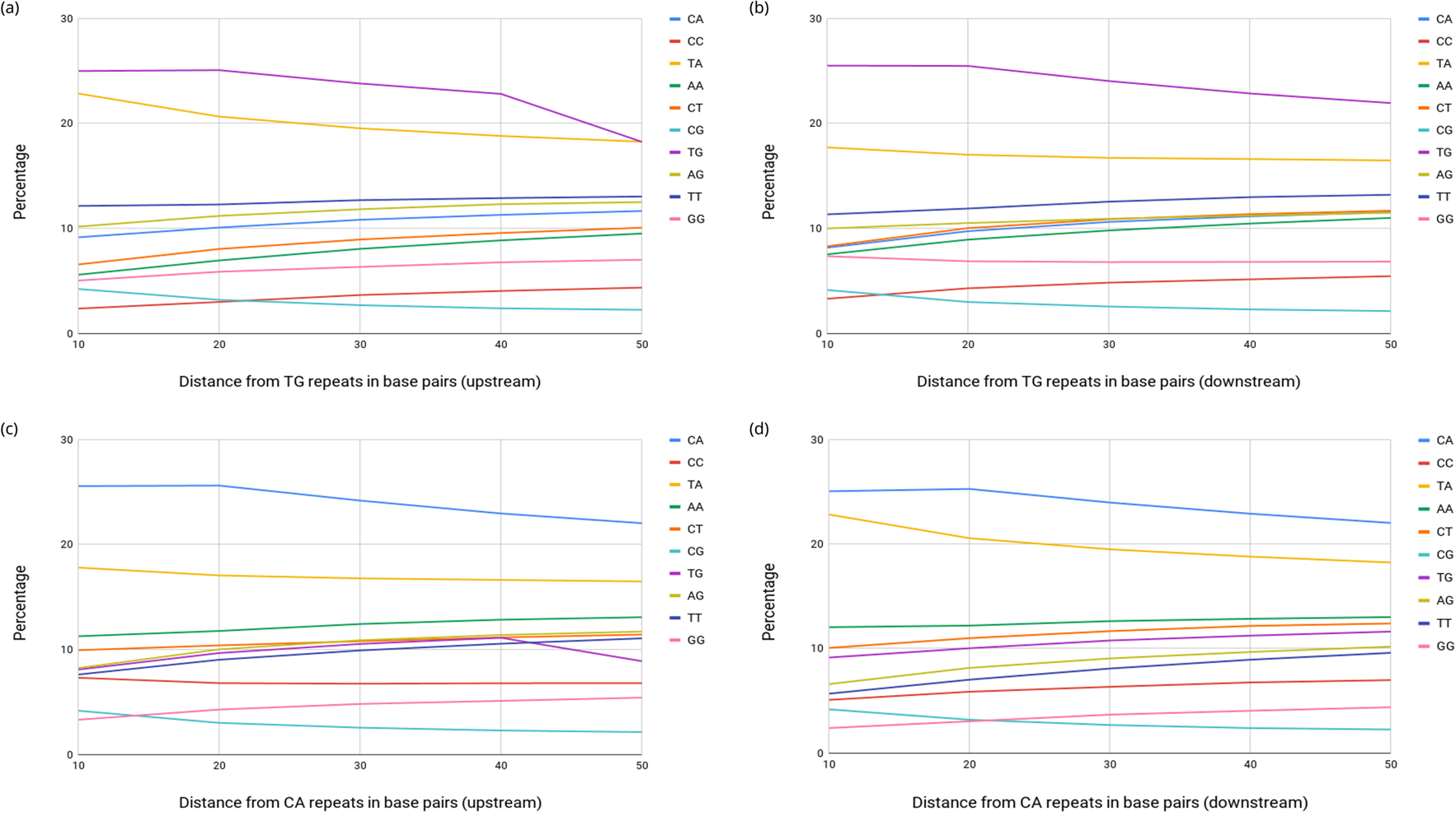
Dinucleotide composition of the 50 bp flanking region upstream and downstream of TG/CA repeats in human.

### 2.7. The genomic alignments from UCSC browser

Most of the analysis and results shown in this study are from the pairwise alignments of the human genome with other primate genomes. Therefore, to illustrate the observations and results presented in the above sections, some examples of alignments including humans and all sixteen primate genomes together were retrieved from the UCSC Genome Browser [30] and shown in Figure S6 (A-E). The observations from the screenshots of selected cases to highlight the different types of repeat conversions, such as elongation of TG/CA repeats as compared to other primates (Case B), point mutations leading to conversion of different repeat types to TG/CA repeats (Case E) and their expansions further strengthen the results of this study.

## 3. Discussion

TG/CA microsatellites are the most abundant dinucleotide repeats in the human genome [4, 8, 12]. They exhibit multifaceted functional roles, such as modulation of gene expression [26], recombination [11] and intron splicing due to their polymorphic nature [27]. Therefore, a comparative study of these repeats in primate genomes, which originated from a common ancestor, is likely to provide useful insights into their conservation and evolution. In this study, sixteen evolutionary distant primate genomes were examined through pairwise alignment with the human genome. The total length of their alignments with the human genome varied; however, this length variation can be attributed to the evolutionary distance between human and other primate species. Therefore, for the comparative analysis, normalized proportions were used in all calculations to minimize these variations.

The comparative analysis of (TG/CA)_n_ repeats at orthologous locations in the pairwise alignments of the human and other primate genomes shows that the total number of these repeats in human is higher than the other primates. The incidence of these orthologous repeats in the aligned regions decreases with increasing evolutionary distance from human to bushbaby, yet it is interesting to note that more than 50% of these repeats can still be found at the orthologous locations in bushbaby, which diverged from human about 74 million years ago. In a few cases (Figure S6, Case D), though chimpanzee genome is the closest to the human genome, some of the human (TG/CA)_n_ repeats were found in a mutated form in more distant primates but were absent in chimpanzees. It seems less likely that this could be due to the gaps in the sequencing of the chimpanzee genome since the regions were present in other more distant primates and, for those primates, the sequencing is lagging far behind than chimpanzee genome sequencing. Therefore, it indicates that some of these repeat units could be lost or added during the evolution. All primates originated from a common primordial ancestor for whom the number and location of these repeats cannot be known. Therefore, it can be argued that those repeats which are present in human and also present in any of the distant primate might have remained conserved at that location (allowing length variation) and could be originally present in the common ancestor, because it is less likely that a repeat at a particular location could originate independently in the two diverged genomes.

The lengths of the (TG/CA)_n_ repeats were observed to be longer in humans as compared to the other primates (Figure 4). Similar observations were also made in a few previous studies using small genomic regions or genetic markers from human and chimpanzee genomes [28, 29]. The genome-wide analysis and comparison of sixteen primates to humans, spanning an evolutionary distance of about 74 million years conducted here confirms that these repeats have increased in length in the human genome. An example of HSD11B2 gene where CA repeats are present in intron-1 (Figure S6, Case F) shows that these repeats have elongated in human as compared to other primates (except Orangutan). HSD11B2 gene encodes for an isozyme of 11β-hydroxysteroid dehydrogenase associated with mineralocorticoid regulation in kidney. Higher activity of HSD11B2 gene lowers mineralocorticoid levels, thus preventing familial hypertension [31]. Previous studies have shown direct association between HSD11B2 expression and length of CA repeat in its first intron [32]. This example suggests that selection and elongation of CA repeats in human HSD11B2 intronic region as compared to other primates has an advantage with respect to the regulation of gene expression. Previous studies have also shown role of TG/CA repeats in transcription regulation in genes like VLDLR, GHR [33, 34] etc. Further, the observed the elongation of medium and short repeats in this study indicates that such repeats have greater tendency to expand as compared to small repeats. This also corroborates with previous studies which suggested that the slippage rate is correlated to short tandem repeat (STR) length [35, 36, 37, 38].

Since it is well known that (TG/CA)_n_ repeats exhibit length polymorphism between different individuals of the same species, the question emerges whether the increase in repeat length is due to elongation during evolution or due to common polymorphism. Our study shows that both the length and number of these repeats have increased gradually over evolutionary time, thus supporting the notion that these repeats are longer in humans as compared to the other primates mainly due to elongation. Additionally, the analysis of orthologous regions in the primate alignments shows that at several loci where a (TG/CA)_n_ repeat exists in the human genome, the corresponding primate regions have either a mutated version of these repeats, or they have a different dinucleotide repeat (mainly TA and GA). This further supports the above notion that it is not only polymorphism but conversion of other dinucleotide repeats to (TG/CA)_n_ repeats happened during evolution, pointing towards a positive selection of these repeats in the human genome.

It was also observed that at several loci where a perfect (TG/CA)_n_ repeat occurs in the human genome the corresponding repeat in the primate genome had one or more point mutations within the repeat stretch. Along with the previous finding that the total number of these repeats is the highest in the human genome, this observation further points toward their positive selection in the human genome. The frequencies of transitions, A->G and C->T in TG repeat regions and T->C and G->A in CA repeat regions, are much higher than the genome-wide transition frequencies. These nucleotide transitions, at the dinucleotide level (mainly TA->TG and TA-> CA), also increase the probability of their conversion to TG or CA repeats.

In most cases these conversions occur in single step increments (TA)_1_→((TG)_1_ or (CA)_1_), but multi-step conversions of as many as 25 units have also been observed (Supplementary table S1, Supplementary table S2). Such multi-step conversions appear to be a recent phenomenon, as it is observed more often in the alignments where the primates are more closely related to humans. However, the few-step conversions were observed in all non-human primates. These observations can be interpreted in either of two ways: the corresponding imperfect repeats in the non-human primate have become perfect in human, or the corresponding perfect repeats in human have become imperfect in the non-human primate. If the former argument is true, it supports the hypothesis that these repeats are evolutionarily favored in humans. In support of this argument, it was observed that the step conversion frequency increases with increasing evolutionary distance (Supplementary table S1, Supplementary table S2). Chimpanzee, being the closest to humans, has the most similar number of repeats and the least number of mutated residues in the repeat stretches as human as compared to the other primates. In contrast, the most distal primate bushbaby showed a higher number of mutations in the repeat stretches. Taken together, this implies purifying selection within the repeat regions towards the formation of perfect (TG/CA)_n_ repeat stretches in humans.

The flanking regions of (TG/CA)_n_ repeats might play a role in their expansion. It was observed that the most adjacent flanking regions of the repeats are either rich in degenerated or mutated short (TG/CA)_n_ repeats, or TG/CA dinucleotides, or other dinucleotides containing those nucleotides which have higher transition or transversion frequencies for forming the respective TG or CA unit of the repeat. However, these remnant nucleotides flanking the (TG/CA)_n_ repeats could also be the result of random mutations in perfect repeat stretches [6, 7]. Another possibility, as suggested in other studies, could be that these repeats influence the flanking regions, induce local mutational biases and convergent evolution around microsatellites [39, 40]. Also, since the neighboring sequences of SSRs (Simple Sequence Repeats) typically contain degenerated motifs that result from point mutations on the edge, it might play a key role in stabilizing the SSR. This indicates the role of these repeats in the evolution of the flanking regions. However, since the composition of the flanking regions is very similar in all primate genomes, it suggests that the flanking region plays a role in length polymorphism and repeat length dynamics due to random mutations within individuals of the same primate species.

The previously proposed mechanism of repeat-length polymorphism suggests that length changes mainly arise by polymerase slippage during DNA replication; and due to these slippage mutations, shorter repeats tend to expand and longer repeats tend to contract [5, 6, 7, 9] (Figure 8a). This satisfactorily explains the repeat-length polymorphisms observed among individuals of the same species, but it does not explain the selection and expansion of these repeats from distant primates to humans. Thus, from observations made in this study, we propose that directional nucleotide transition and transversion also play a role in (TG/CA)_n_ repeat expansion in the human genome (Figure 8). All four mechanisms shown in Figure 8 appear to play a role in repeat elongation. Figure 8a shows the previously known mechanism of repeat elongation and contraction within individuals of the same species. Figure 8b shows the directional transition and transversion from primates to humans proposed here to play a role in repeat elongation in the human genome. Due to these transitions and transversions, the other dinucleotide repeats (such as TA and CG) present in other primate genomes at locations orthologous to the human (TG/CA)_n_ repeats, over a period of time, were converted to TG or CA repeats (Figure 8c). The flanking regions, both upstream and downstream of (TG/CA)_n_ repeats, which are rich in those dinucleotides having a higher frequency of conversion to (TG/CA)_n_ repeats, also appear to play a role in the elongation of these repeats in the human genome (Figure 8d).

**Figure 8.**
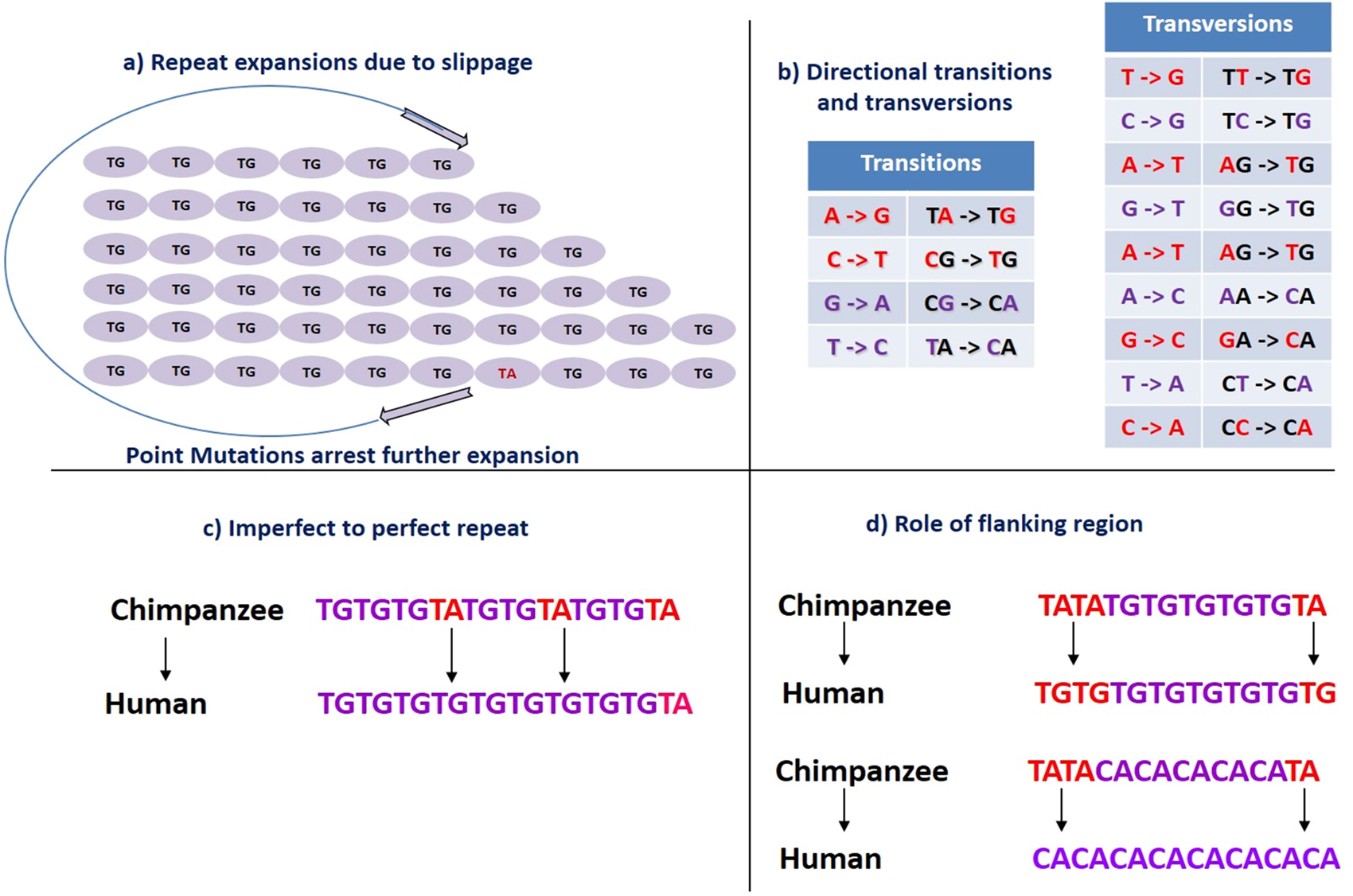
Model for directional selection of (TG/CA)n repeats in the human genome

There appears to be a positive selection or preference for (TG/CA)_n_ repeats over other dinucleotide repeats in the human genome. The conversion of a few units of TA to TG or CA seems plausible, but the conversion of long TA repeat regions (≥20 units) to perfect TG or CA repeat regions (Figure S6, Case E) is incredible, yet occurs, strongly suggesting a positive selection for these repeats. It would be interesting to further study the significance of such long (TG/CA)_n_ repeat regions in the human genome which originated from the conversion of long TA or other dinucleotide repeats found in primates. The abundance and selection of (TG/CA)_n_ repeats may be attributed to the multifaceted functional roles of these repeats in the human genome. As (TG/CA)_n_ repeats show high propensity to form Z-DNA, evolution and elongation of such repeats in the human genome could help in heightened degree of regulation of gene expression.

Using pairwise alignments instead of aligning all genomes together is a limitation of the current study since at the time when the data was retrieved for the current study, only the pairwise alignments were available. Therefore, we could not perform the analysis using all genomes together. However, to support this study, we have presented alignments from a few selected repeat regions which were present at orthologous locations in all or most genomes (Figure S6).

Taken together, this study provides clues on the selection and expansion of TG/CA repeats in the human genome for their vital role in recombination, alternate splicing and gene regulation. The results from the study provides leads for further studies by using the multiple sequence alignments from more primate genomes in a phylogenetic framework to provide insights into the role of these repeats in primate evolution, particularly in the humans where they show the highest abundance.

## 4. Methods

### 4.1. Retrieval of sequence alignments

Pairwise chromosomal alignments of the primates used in this study (as per Table 1) with the human genome assembly (build hg38) were retrieved from UCSC genome browser (https://hgdownload.soe.ucsc.edu/downloads.html#human) in axtNet alignment format. These alignments were carried out using the BLASTZ program. Ideally, a single best orthologous match for each human region matching the corresponding chromosome of the aligned primate genome is expected after alignment. Though the alignment program looks for the best match to each human region, in some alignments one sequence from the human genome may show a match with more than one sequence in the aligned primate genome due to duplications in the primate genome. Thus, for all the primates we’ve analyzed, only those alignments that were ≥ 1 kb and belonged to the same homologous chromosome were selected.

### 4.2. Mapping and characterization of (TG/CA)n repeats

An inhouse Python3 script was used for identification of uninterrupted (TG/CA)n repeats (n≥6 units) in the human and other primate chromosome alignments. (TG/CA)n repeats were grouped into three categories (Types I, II and III), according to their length and biological properties as defined previously. Type I (TG/CA)n repeats, in the range 6 ≤ n < 12 units, are short repeats and are based on the observation that a repeat length of 8 units (n = 8) is the minimum likely to be polymorphic. Type II (TG/CA)n repeats in the range of 12 ≤ n < 23 units, are medium repeats and are based on the observation that more than 93% of the (CA)n repeats of n ≥12 units are polymorphic. They were all counted using another inhouse Python3 script. Furthermore, repeats of this length have also been shown to have preferential binding to nuclear factors as compared to short repeats, and they can also stimulate mRNA splicing. Type III (TG/CA)n repeats ranging from n ≥ 23 units are long repeats and they have a propensity to adopt structures such as Z-DNA. Other studies have shown that (TG/CA)n repeats of length greater than 22.5 units can stimulate recombination. In this work, Type I, II and III repeats are referred to as short, medium and long repeats, respectively.

### 4.3. (TG/CA) repeats in human genes and analysis of the flanking regions

The human gene information (same build as the human genome) was also retrieved from UCSC (more details given in the next section). The repeats were mapped in the intragenic regions (including both introns and exons) and the intergenic regions of the human genes found in the chromosomal alignments. This was done using an in-house Python script.

Using the above mentioned genome assembly and the same algorithm for mapping and characterization for (TG/CA)n repeats, flanking regions in intervals of 10 (up to 50 and overlapping) were identified both upstream and downstream of the repeat region. They were analyzed for the prevalence of single nucleotides and dinucleotides (which had potential for transition or transversion to TG / CA) using an in-house Python Script. Their corresponding percentage with respect to the size of the flanking region being considered was calculated using the same.

### 4.4. Gene data Analysis on Human-Chimpanzee alignment file

All the gene data analysis of TG/CA repeats in introns, exons or on the intron-exon boundary were done on human-chimpanzee alignment file. The data of all the exons were extracted from UCSC Table browser (https://genome.ucsc.edu/cgi-bin/hgTables) by generating a gtf file for the appropriate human genome data (hg38) and using GENCODE as the track for our data of exons and thus our genes. After matching the exon data with the data obtained for TG/CA repeat regions in the alignment file of human-chimpanzee, factors like repeat length, repeat region (exons, introns or intron-exon boundary) and the gene and chromosome information were obtained to visualize what that section of DNA looked like in the other primates. Selecting human-chimpanzee alignment for this analysis, which is closest to human on evolutionary timescale and covers most of the human genome, was necessary to identify recent changes in TG/CA repeat length in different parts of the genome as well as to observe how these repeats changed across the whole primate lineage.

### 4.5. Construction of Evolutionary / Phylogenetic Tree

All the data and the backbone figure was obtained from TimeTree, a public evolutionary distance mapper (http://www.timetree.org/) which compiles information from various studies [41].

### 4.6. Statistical analysis

Pearson’s correlation coefficient was calculated using in-built Microsoft Excel function. Based on the Pearson’s correlation coefficient, t-statistic was obtained using the following formula:

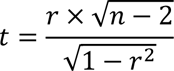

Where, r = Pearson’s correlation coefficient, n = sample size, t = t-statistic to be calculated.

Based on this t-statistic, two-tailed analysis was carried out to obtain corresponding p-value using Microsoft Excel’s in-built function.

## Author’s contributions

VKS conceived of the idea and carried out the entire analysis in seven primate genomes, prepared the figures and wrote the manuscript. Later, SM, ASM and PH repeated the genomic analysis on a larger set of 16 primate genomes and prepared the final figures and tables. VKS, ASM and SM prepared the final manuscript.

## Acknowledgements

We thank IISER Bhopal for intramural funding. Mr. Aditya Malwe thank IISER Bhopal for the fellowship.

## Compliance with ethical standards

Not Applicable

## Supplementary information

**Table S1.** Information about primate species used for the analysis

**Table S2.** Step conversion frequency normalized by the genome size (as available in genomic alignments) for other dinucleotide repeats from non-human primate genomes that got converted to TG repeats in human genome

**Table S3.** Step conversion frequency normalized by the genome size (as available in alignments) for other dinucleotide repeats in non-human primate genomes that got converted to CA repeats in the human genome

**Figure S1.** Overview of (TG/CA)_n_ repeat dynamics in primate genomes

**Figure S2:** Percentage of human genome covered in alignment with other primates.

**Figure S3.** Correlation between evolutionary distance and repeat elongation, contraction and conservation in human as compared to primates

**Figure S4.** Distribution and number of contracted, equal-length and elongated repeats in different genomic regions.

**Figure S5.** Genome-wide frequency of transitions and transversions in the human and other primate genomic alignments

**Figure S6.** Selected examples of repeat length dynamics in primate genomes

## References

1 Dib C, Fauré S, Fizames C, Samson D, Drouot N et al. (1996) A comprehensive genetic map of the human genome based on 5,264 microsatellites. Nature. 380:152–154.

2 Kong A, Gudbjartsson DF, Sainz J, Jonsdottir GM, Gudjonsson SA, et al. (2002) A high-resolution recombination map of the human genome. Nat.Genet. 31:241–247.

3 Ota T and Kimura M. (1973) A model of mutation appropriate to estimate the number of electrophoretically detectable alleles in a finite population. Genet.Res. 22:201–204.

4 Ellegren, H. (2004) Microsatellites: simple sequences with complex evolution. Nat.Rev.Genet. 5:435–445.

5 Kruglyak S, Durrett RT, Schug MD, and Aquadro CF. (1998) Equilibrium distributions of microsatellite repeat length resulting from a balance between slippage events and point mutations. Proc.Natl.Acad.Sci.U.S.A. 95:10774–10778.

6 Levinson G and Gutman GA. (1987) Slipped-strand mispairing: a major mechanism for DNA sequence evolution. Mol.Biol.Evol. 4:203–221.

7 Strand M, Prolla TA, Liskay RM, and Petes TD. (1993) Destabilization of tracts of simple repetitive DNA in yeast by mutations affecting DNA mismatch repair. Nature. 365:274–276.

8 Lander ES, Linton LM, Birren B, Nusbaum C, Zody MC, et al. (2001) Initial sequencing and analysis of the human genome. Nature. 409:860–921.

9 Rose O and Falush D. (1998) A threshold size for microsatellite expansion. Mol.Biol.Evol. 15:613–615.

10 Leclercq S, Rivals E, and Jarne P. (2010) DNA slippage occurs at microsatellite loci without minimal threshold length in humans: a comparative genomic approach. Genome Biol.Evol. 2:325–35.

11 Majewski J and Ott J. (2000) GT repeats are associated with recombination on human chromosome 22. Genome Res. 10:1108–1114.

12 Sharma VK, Brahmachari SK, and Ramachandran S. (2005) (TG/CA)n repeats in human gene families: abundance and selective patterns of distribution according to function and gene length. BMC.Genomics. 6:83.

13 Tautz D and Renz M. (1984) Simple sequences are ubiquitous repetitive components of eukaryotic genomes. Nucleic Acids Res. 12:4127–4138.

14 Schlotterer C and Tautz D. (1992) Slippage synthesis of simple sequence DNA. Nucleic Acids Res. 20:211–215.

15 Richard GF and Paques F. (2000) Mini- and microsatellite expansions: the recombination connection. EMBO Rep. 1:122–126.

16 Meera G, Ramesh N, and Brahmachari SK. (1989) Zintrons in rat alpha-lactalbumin gene. FEBS Lett. 251:245–249.

17 Naylor LH and Clark EM. (1990) d(TG)n.d(CA)n sequences upstream of the rat prolactin gene form Z-DNA and inhibit gene transcription. Nucleic Acids Res. 18:1595–1601.

18 Nordheim A and Rich A. (1983) The sequence (dC-dA)n X (dG-dT)n forms left-handed Z-DNA in negatively supercoiled plasmids. Proc.Natl.Acad.Sci.U.S.A. 80:1821–1825.

19 Wang, G., & Vasquez, K. M. (2007). Z-DNA, an active element in the genome. Front Biosci, 12(4424), 38.

20 Oh, D. B., Kim, Y. G., & Rich, A. (2002). Z-DNA-binding proteins can act as potent effectors of gene expression in vivo. Proceedings of the National Academy of Sciences, 99(26), 16666–16671.

21 Pravica V, Asderakis A, Perrey C, Hajeer A, Sinnott PJ, et al. (1999) In vitro production of IFN-gamma correlates with CA repeat polymorphism in the human IFN-gamma gene. Eur.J.Immunogenet. 26:1–3.

22 Shimajiri S, Arima N, Tanimoto A, Murata Y, Hamada T, et al. (1999) Shortened microsatellite d(CA)21 sequence down-regulates promoter activity of matrix metalloproteinase 9 gene. FEBS Lett. 455:70–74.

23 Sharma VK, Rao C, Sharma A, Brahmachari SK, and Ramachandran S. (2003) (TG:CA)(n) repeats in human housekeeping genes. J.Biomol.Struct.Dyn. 21:303–310.

24 Sharma VK, Sharma A, Kumar N, Khandelwal M, Mandapati KK, et al. (2006) Expoldb: expression linked polymorphism database with inbuilt tools for analysis of expression and simple repeats. BMC.Genomics. 7:258.:258.

25 Sharma VK, Kumar N, Brahmachari SK, and Ramachandran S. (2007) Abundance of dinucleotide repeats and gene expression are inversely correlated: a role for gene function in addition to intron length. Physiol Genomics. 31:96–103.

26 Zheng, J., Xu, H., & Cao, H. (2019). A Long Polymorphic GT Microsatellite within a Gene Promoter Mediates Non-Imprinted Allele-Specific DNA Methylation of a CpG Island in a Goldfish Inter-Strain Hybrid. International journal of molecular sciences, 20(16), 3923.

27 Hui J, Stangl K, Lane WS, and Bindereif A. (2003) HnRNP L stimulates splicing of the eNOS gene by binding to variable-length CA repeats. Nat.Struct.Biol. 10:33–37.

28 Cooper G, Rubinsztein DC, and Amos W (1998) Ascertainment bias cannot entirely account for human microsatellites being longer than their chimpanzee homologues. Hum.Mol.Genet. 7:1425–1429.

29 Webster MT, Smith NG, and Ellegren H. (2002) Microsatellite evolution inferred from human-chimpanzee genomic sequence alignments. Proc.Natl.Acad.Sci.U.S.A. 99:8748–8753.

30 Karolchik D, Barber GP, Casper J, Clawson H, Cline MS, et al. (2014) The UCSC Genome Browser database: update 2014. Nucleic Acids Res. 2014 Jan;42(Database issue):D764–70. doi: 10.1093/nar/gkt1168.

31 Quinkler, M., & Stewart, P. M. (2003). Hypertension and the cortisol-cortisone shuttle. The Journal of Clinical Endocrinology & Metabolism, 88(6), 2384–2392.

32 Agarwal, A. K., Giacchetti, G., Lavery, G., Nikkila, H., Palermo, M., Ricketts, M., … & Stewart, P. M. (2000). CA-Repeat polymorphism in intron 1 of HSD11B2: effects on gene expression and salt sensitivity. Hypertension, 36(2), 187–194.

33 Zhang, W., He, L., Liu, W., Sun, C., & Ratain, M. J. (2009). Exploring the relationship between polymorphic (TG/CA) n repeats in intron 1 regions and gene expression. Human genomics, 3(3), 1–10.

34 Dias, C., Giordano, M., Frechette, R., Bellone, S., Polychronakos, C., Legault, L., … & Goodyer, C. G. (2017). Genetic variations at the human growth hormone receptor (GHR) gene locus are associated with idiopathic short stature. Journal of Cellular and Molecular Medicine, 21(11), 2985–2999.

35 Primmer CR and Ellegren H. (1998) Patterns of molecular evolution in avian microsatellites. Mol.Biol.Evol. 15:997–1008.

36 Sibly RM, Meade A, Boxall N, Wilkinson MJ, Corne DW, et al. (2003) The structure of interrupted human AC microsatellites. Mol.Biol.Evol. 20:453–459.

37 Whittaker JC, Harbord RM, Boxall N, Mackay I, Dawson G, et al. (2003) Likelihood-based estimation of microsatellite mutation rates. Genetics. 164:781–787.

38 Sainudiin R, Durrett RT, Aquadro CF, and Nielsen R. (2004) Microsatellite mutation models: insights from a comparison of humans and chimpanzees. Genetics. 168:383–395.

39 Timsit Y. (1999) DNA structure and polymerase fidelity. J.Mol.Biol. 293:835–853.

40 Vowles EJ and Amos W. (2004) Evidence for widespread convergent evolution around human microsatellites. PLoS.Biol. 2(8): e199.

41 S. Kumar, G. Stecher, M. Suleski, and S.B. Hedges, 2017. TimeTree: a resource for timelines, timetrees, and divergence times. Molecular Biology and Evolution 34: 1812–1819, DOI: 10.1093/molbev/msx116.

42 Schwartz S, Kent WJ, Smit A, Zhang Z, Baertsch R, et al. (2003) Human-mouse alignments with BLASTZ. Genome Res. 13:103–107.

43 Kent WJ, Baertsch R, Hinrichs A, Miller W, and Haussler D. (2003) Evolution’s cauldron: duplication, deletion, and rearrangement in the mouse and human genomes. Proc.Natl.Acad.Sci.U.S.A. 100:11484–11489.

44 Rockman MV and Wray GA. (2002) Abundant raw material for cis-regulatory evolution in humans. Mol.Biol.Evol. 19:1991–2004.

45 Epplen JT, Kyas A, and Maueler W. (1996) Genomic simple repetitive DNAs are targets for differential binding of nuclear proteins. FEBS Lett. 389:92–95.

46 Brahmachari SK, Meera G, Sarkar PS, Balagurumoorthy P, Tripathi J, et al. (1995) Simple repetitive sequences in the genome: structure and functional significance. Electrophoresis. 16:1705–1714.

