## Supplementary figures for "Comparative Analysis of TG/CA Repeats in Sixteen Primate Genomes Reveals the Dynamics and Role of TG/CA Repeats in the Human Genome"

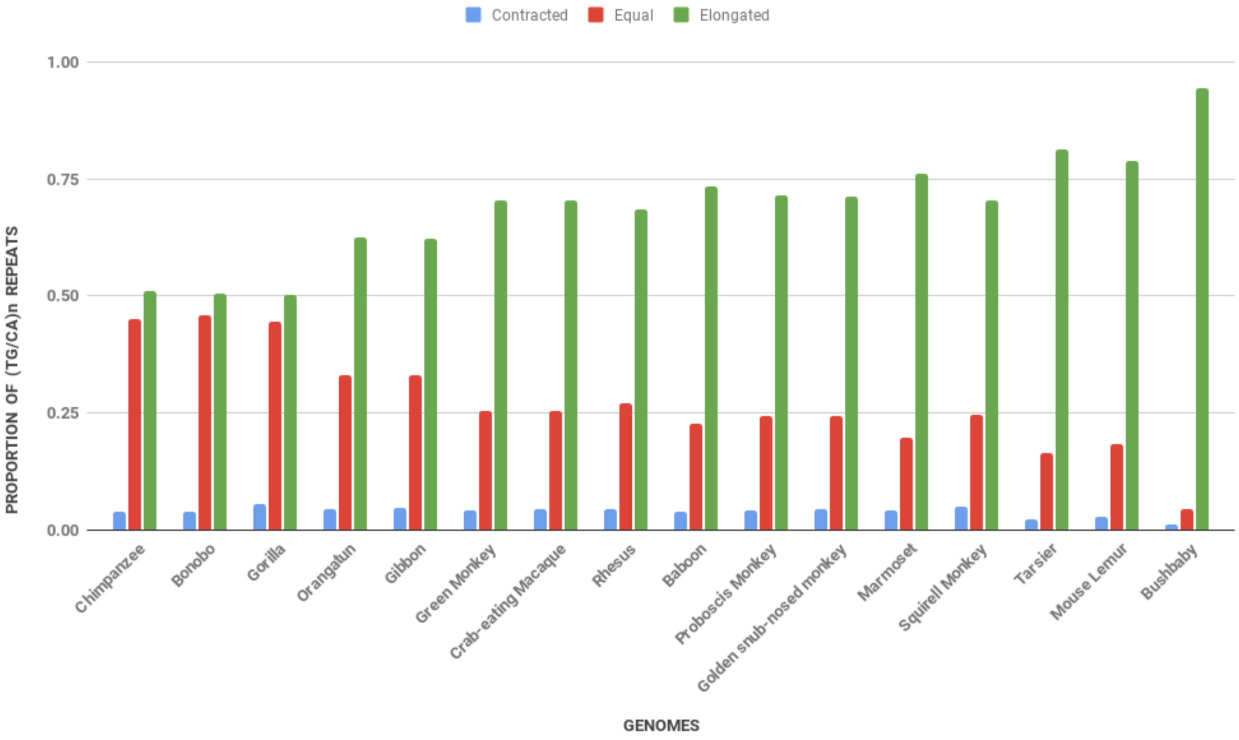


Figure S1: Overview of (TG/CA)n repeat dynamics in primate genomes

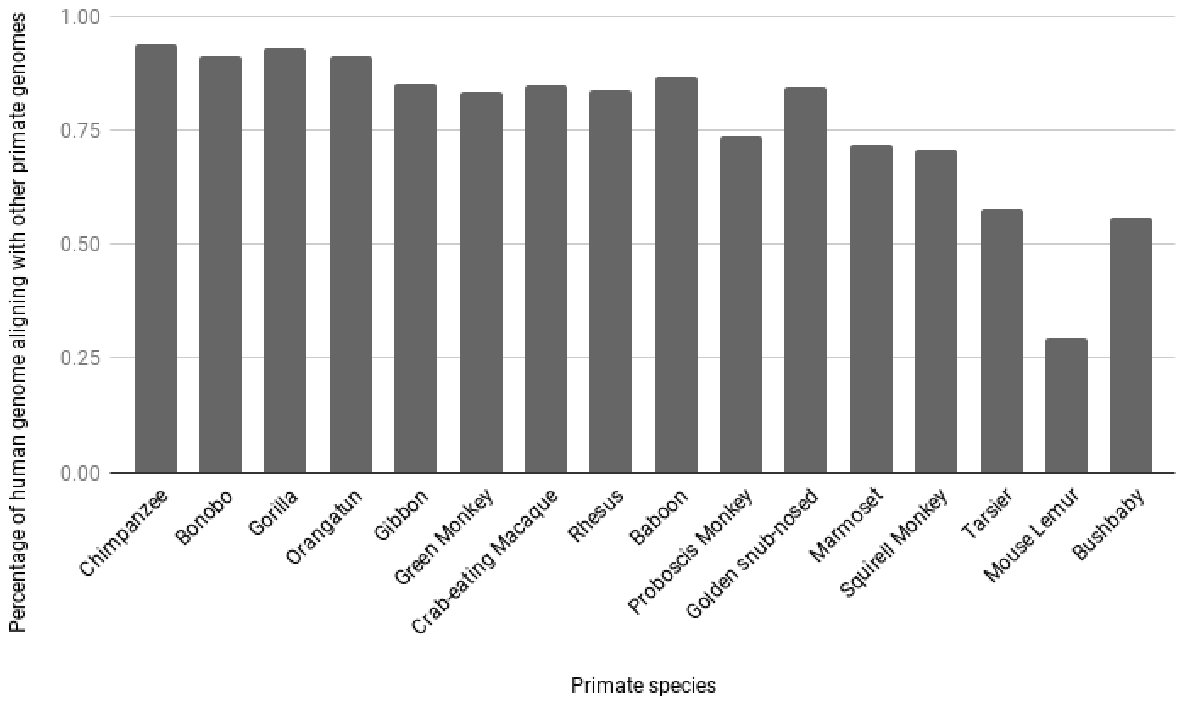


Figure S2: Percentage of human genome covered in alignment with other primates.

Total size of human genome was taken from *https://www.ncbi.nlm.nih.gov/assembly/GCF_009914755.1/#/st*

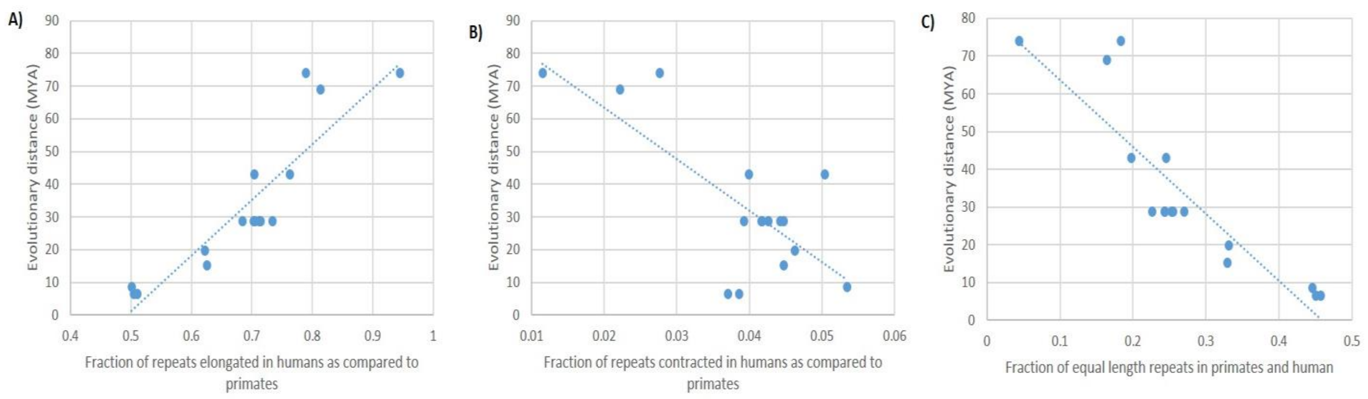


Figure S3: Correlation between evolutionary distance and repeat elongation, contraction and conservation in human as compared to primates

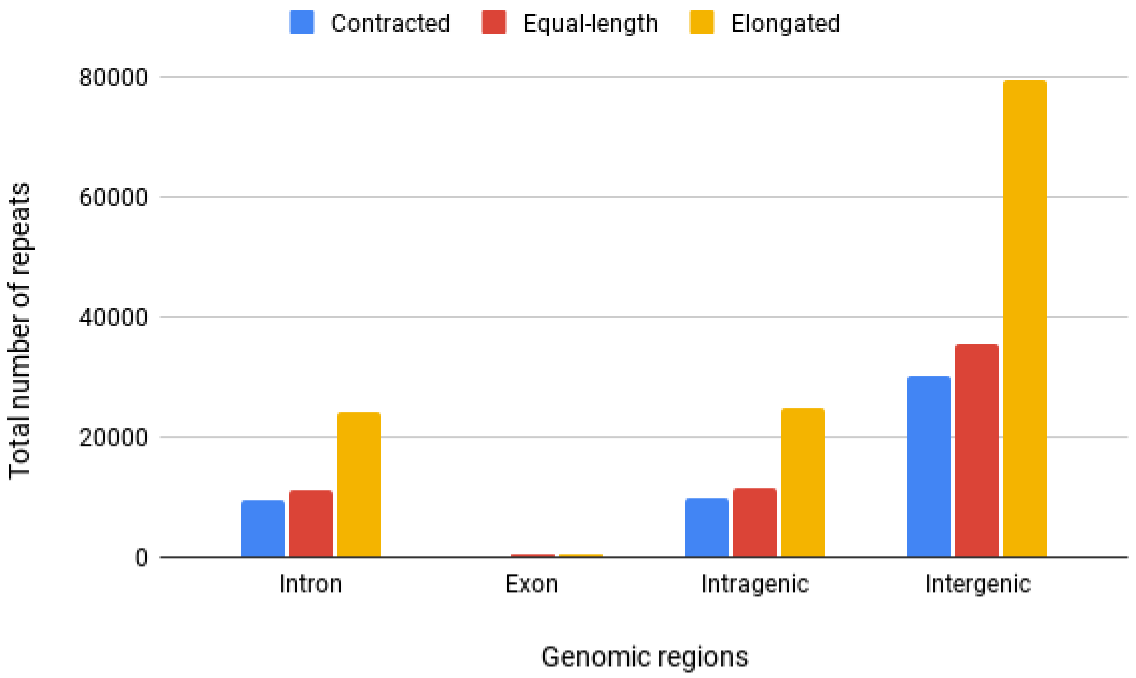


Figure S4: Distribution and number of contracted, equal-length and elongated repeats in different genomic regions.

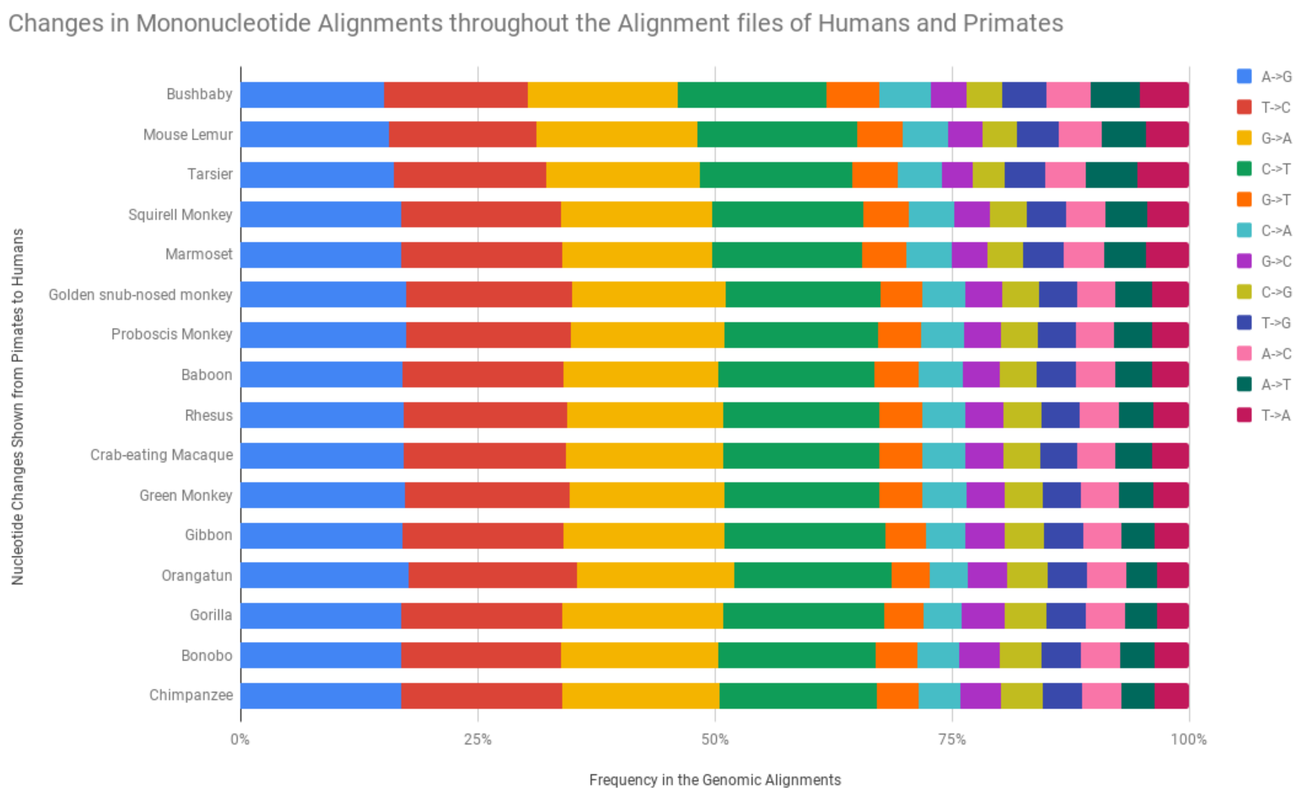


Figure S5: Genome-wide frequency of transitions and transversions in the human and other primate genomic alignments. The arrow on the side of the figure indicates the direction of transition or transversion events from primates to human.

**Figure S6. Selected examples of repeat length dynamics in primate genomes**

In this study, the genomic regions orthologous to human $(TG/CA)_n$ repeats in all other selected primates were compared. To illustrate the observations and results presented in the above sections, some examples have been provided from the aligned regions displayed in the UCSC Genome Browser as shown in cases A to E. Case A shows the length conservation of different legth repeats at orthologous locations in the primate genomes. It was observed that the CA(as well as TG) repeats are highly conserved in human, chimpanzee, gorilla, orangutan, etc. which are closer in the evolutionary tree. However, in the more distant primates (macaque, bushbaby, marmoset, and others), some mutations/omissions were observed in the repeat regions. The next examples (Case B), shows the elongation in repeat length from non-human primates to human and point mutations in orthologous repeats in the primates (with examples showing the extreme cases of elongation from primates to humans) for both TG as well as CA repeats. Case C show greater extent of elongation of TG repeats. Case D shows TG and CA repeats absent in closely related primates like Chimpanzee and Bonobo but present in relatively distant primates like Gorilla and Orangutan. Case E show substitutions leading to formation and elongation of TG/CA repeats. The above cases illustrate the main observations made in this study.

**Case A:** Different length CA repeats conserved across the primate genomes

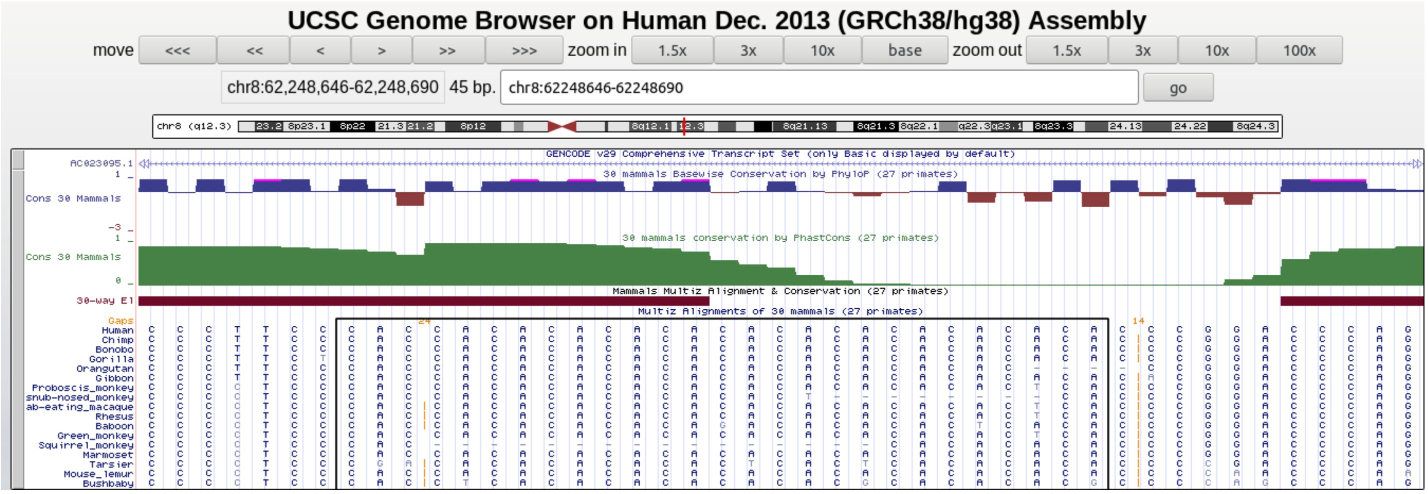

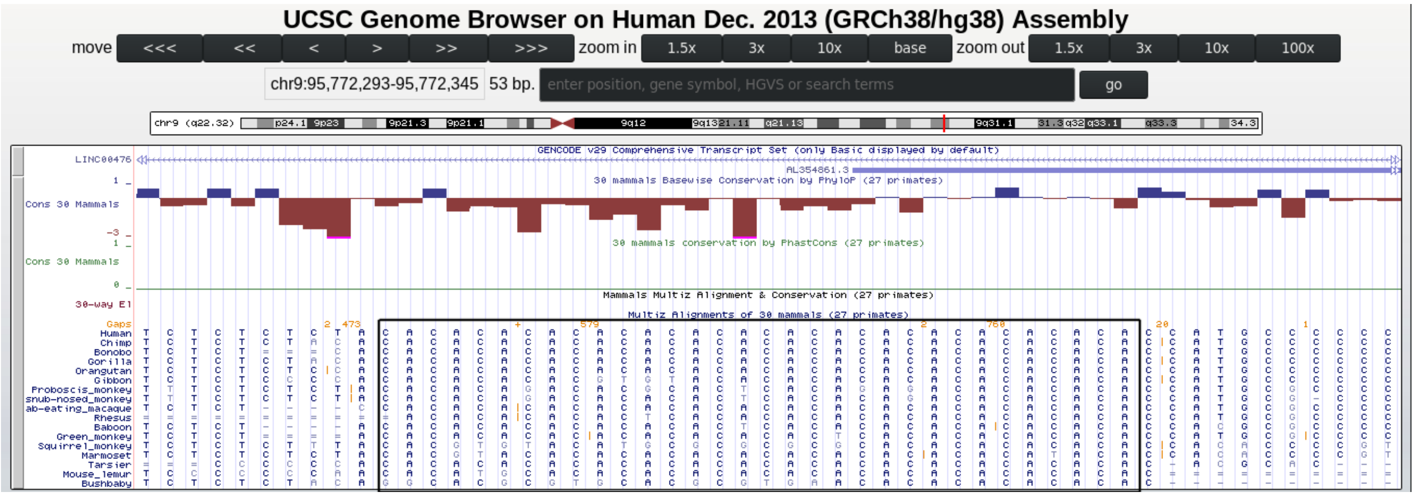


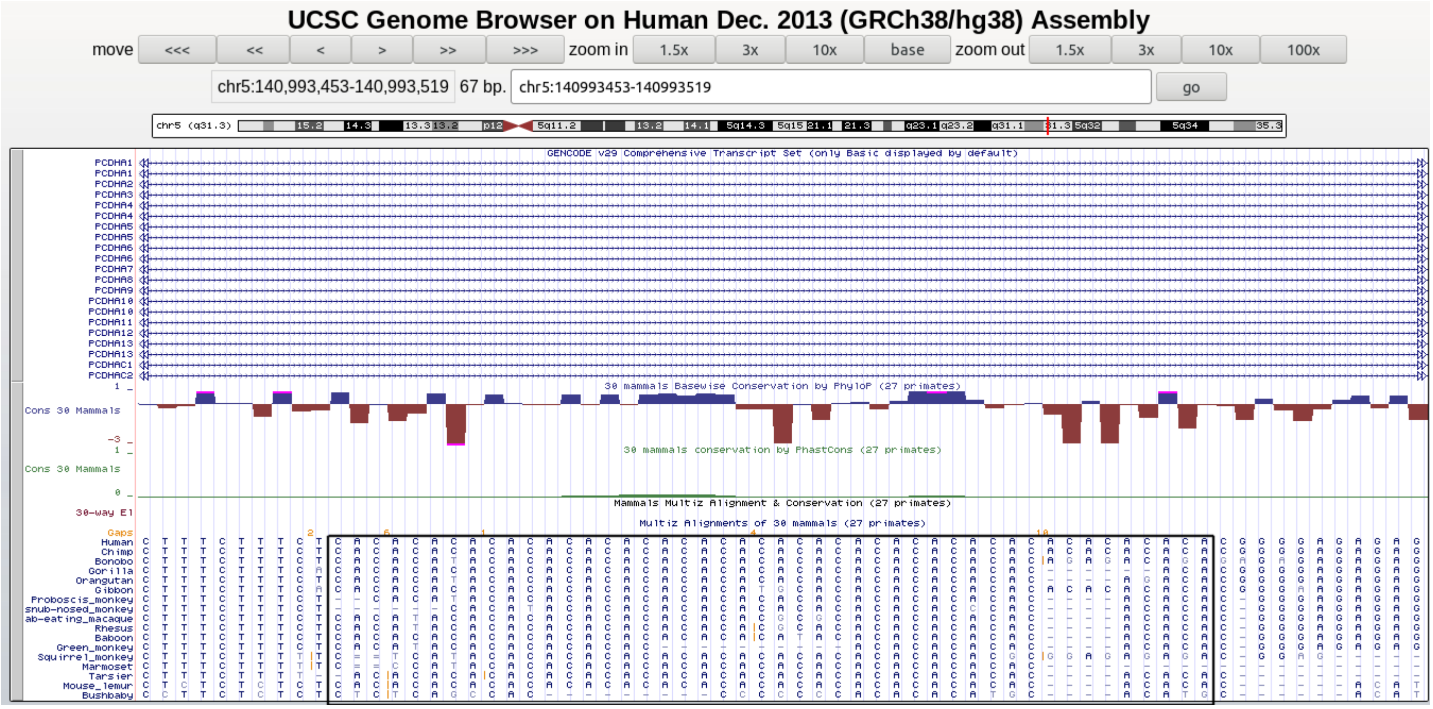


**Case B (a):** Elongation and point mutations in CA repeats from primates to humans

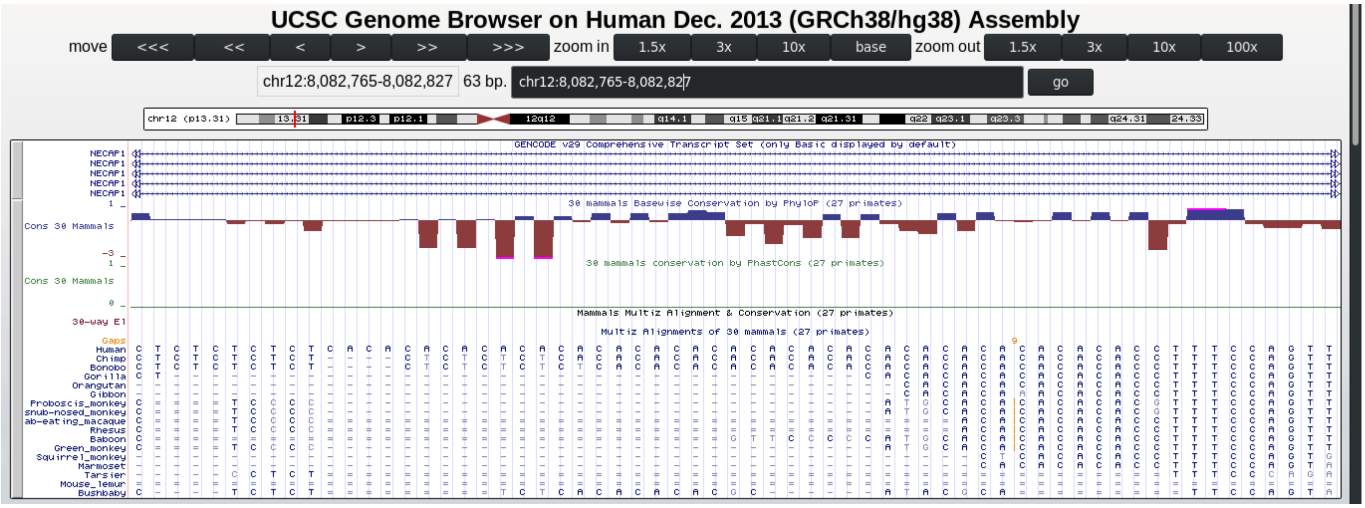


**Case B (b):** Elongation and point mutations in TG repeats from primates to humans

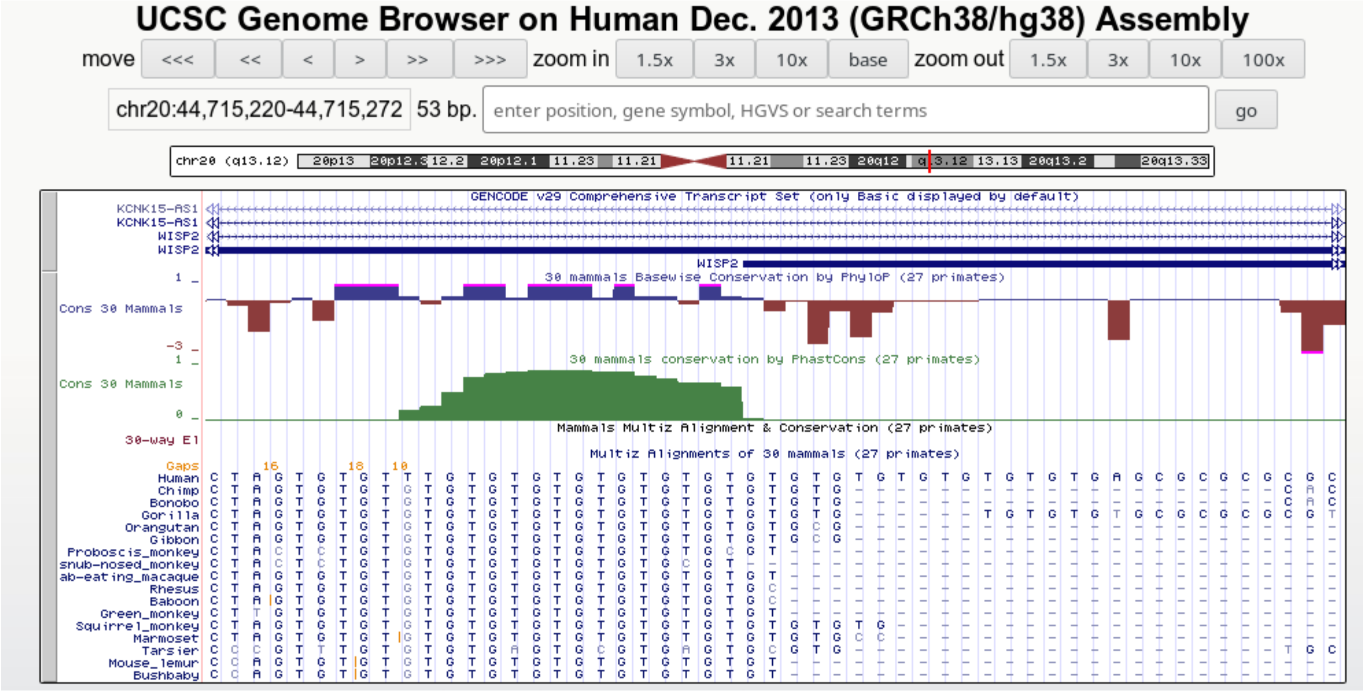

**Case C:** Recently emerged TG repeats elongating from closely related non human primates to human

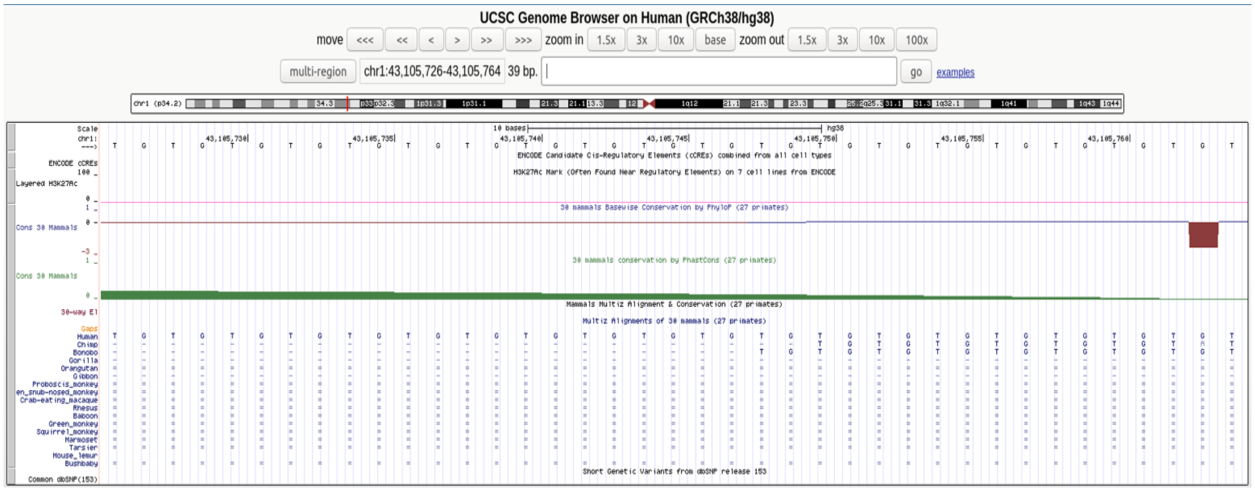


**Case D (a):** TG repeat stretches absent in more closely related primates but present to a limited extent in distantly related primates

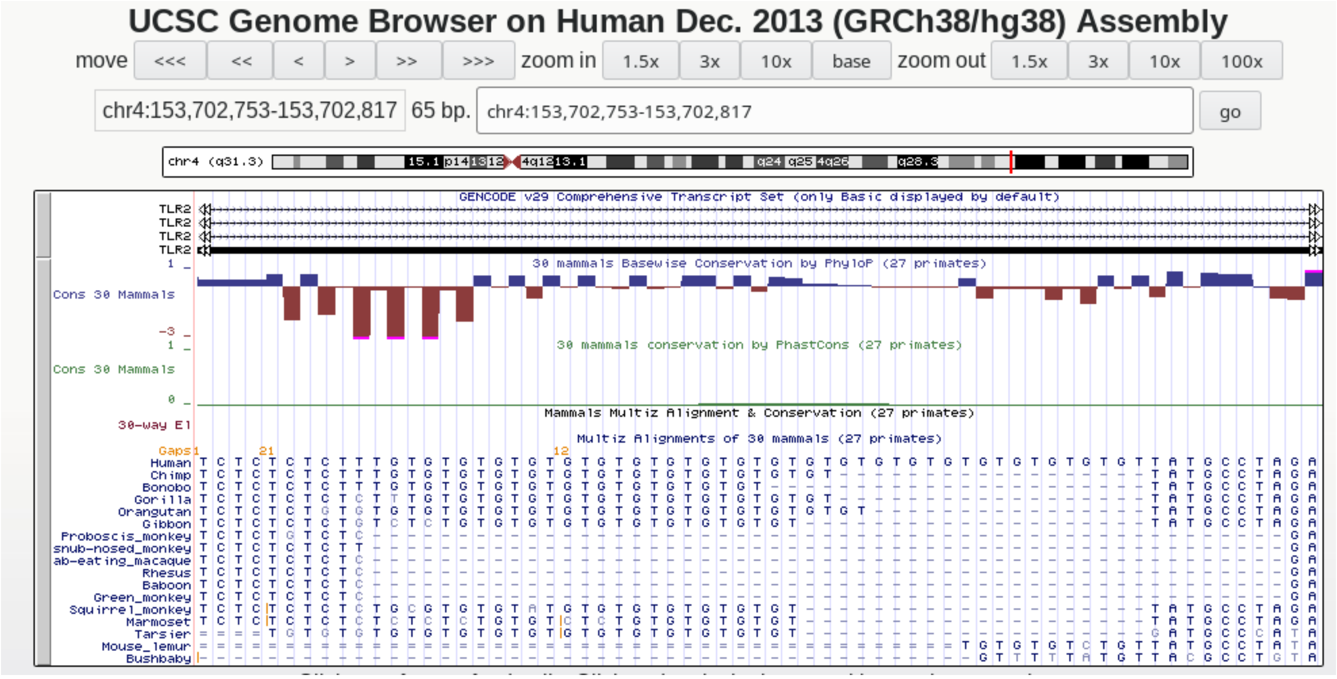


**Case D (b):** CA repeat stretches absent in more closely related primates but present to a limited extent in distantly related primates

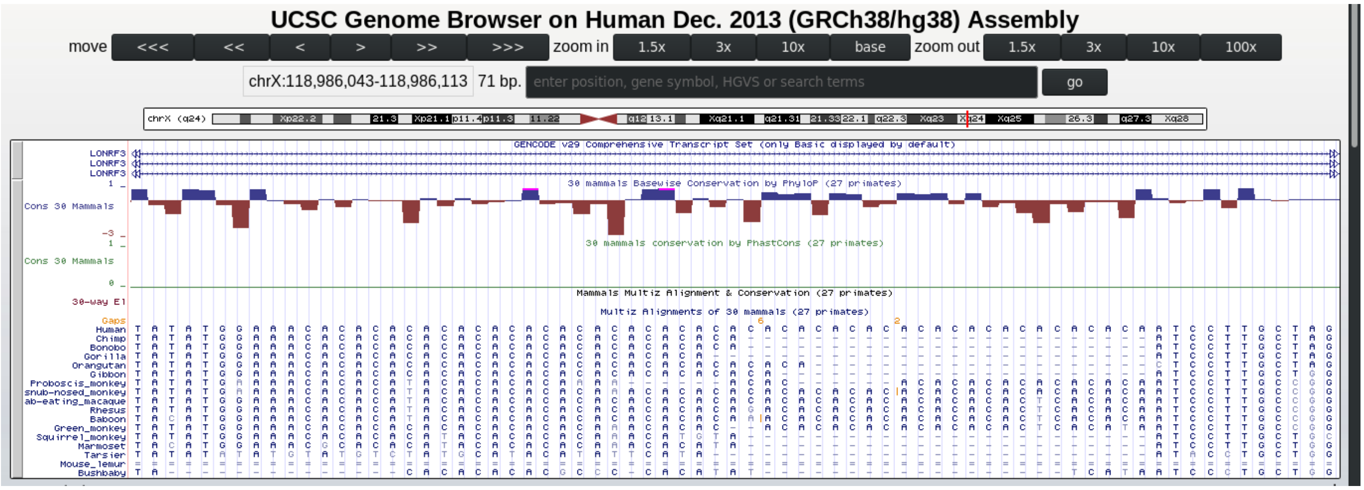


**Case D (c):** Conversion of TA - > TG repeat from non human primates to human that are absent in more closely related primates but incompletely present to a limited extent in distantly related primates

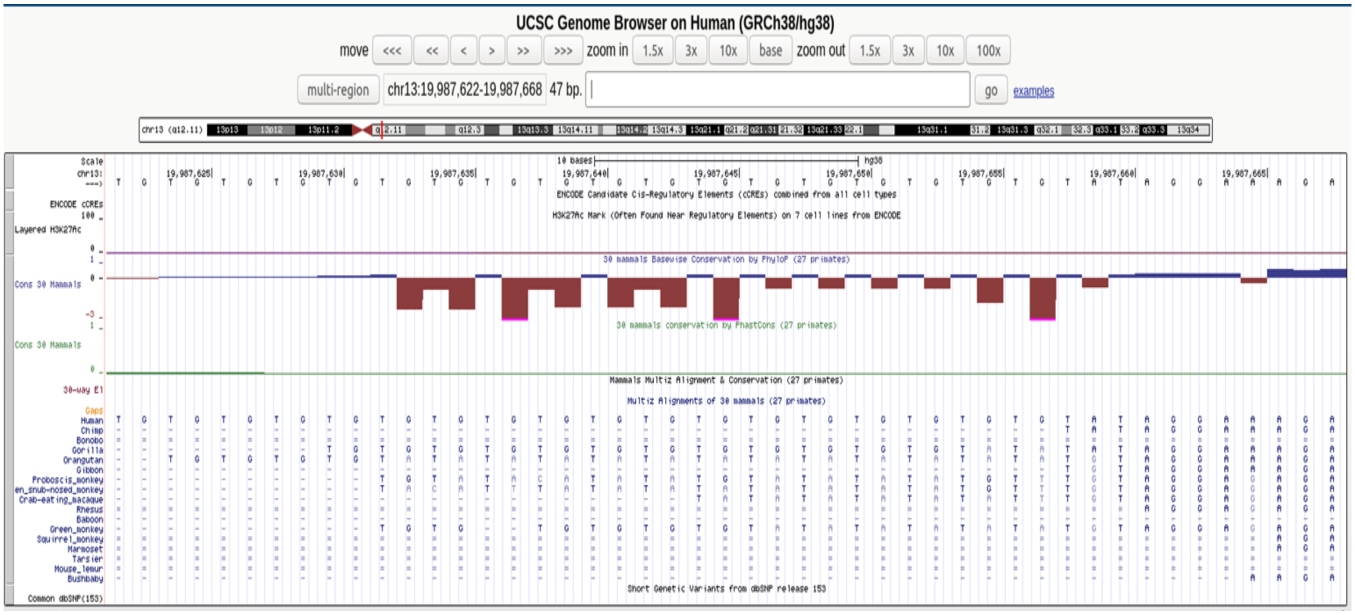

**Case E (a):** Conversion of A->G and TA -> TG from non human primates to human resulting in formation of TG repeats

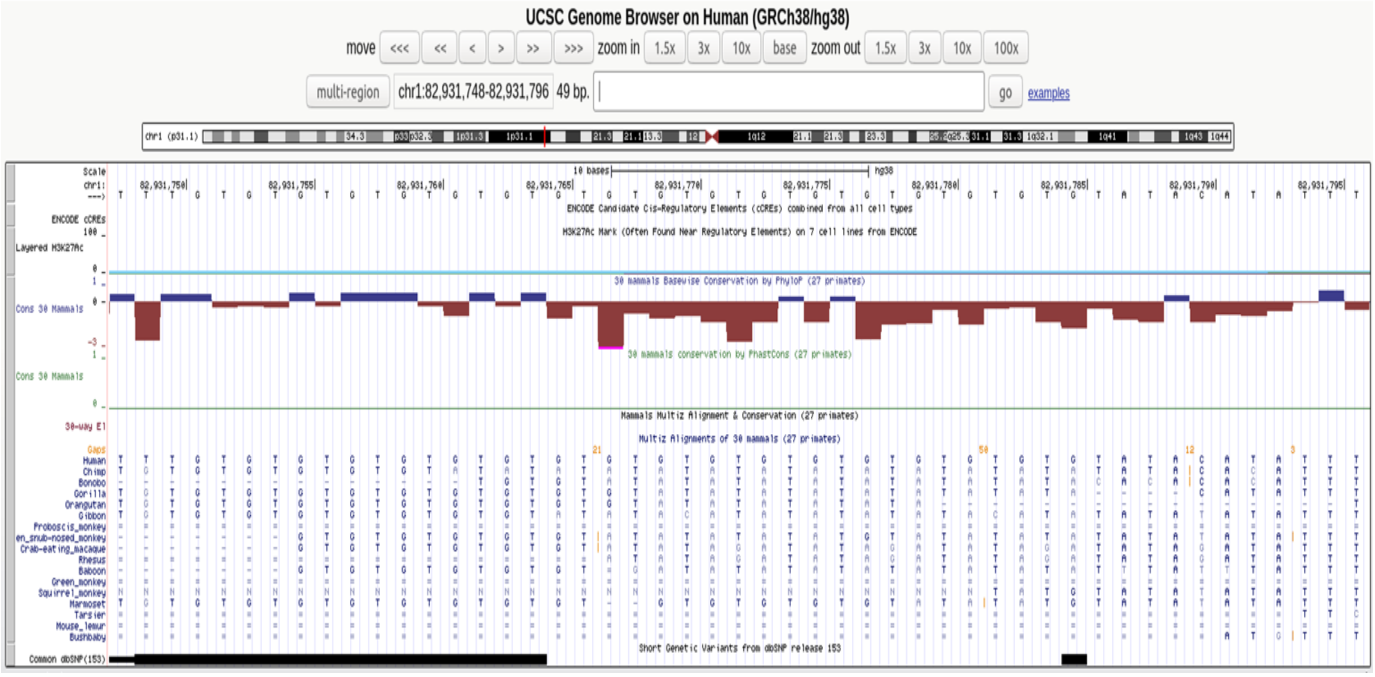


**Case E (b):** Conversion of A->G and TA -> TG from non human primates to human resulting in formation of TG repeats

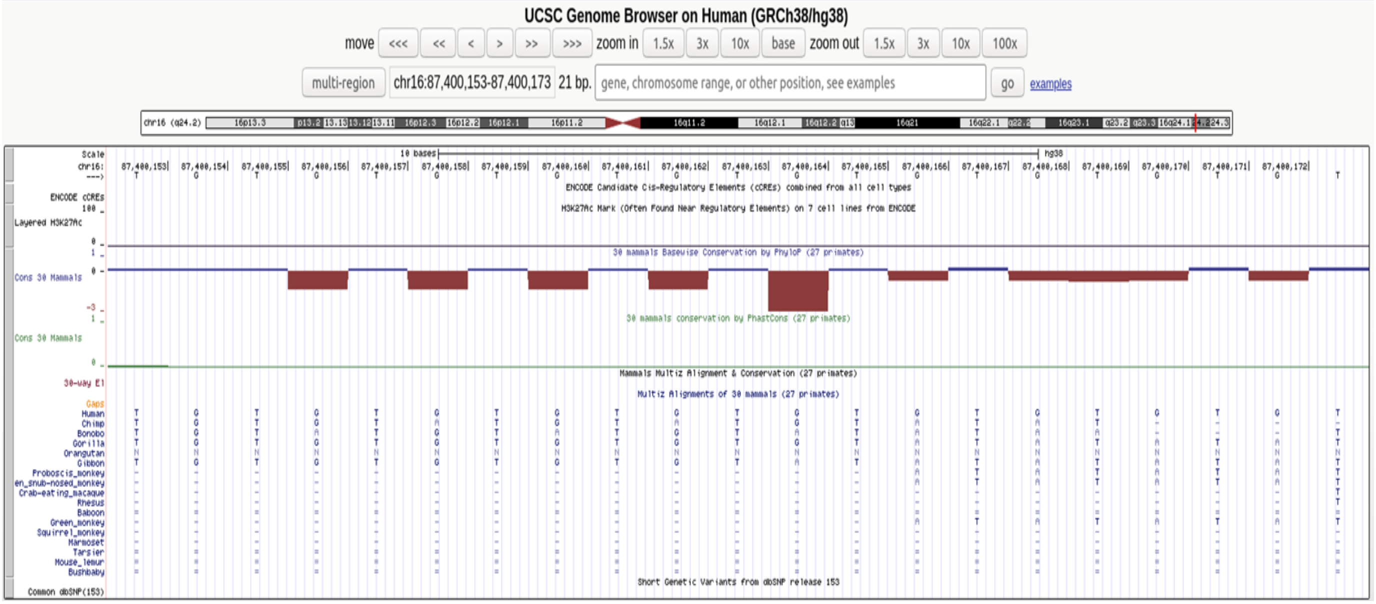


**Case E (c):** Conversion of A->G and TA -> TG resulting in elongation of TG repeats from non human primates to human resulting in formation of TG repeats

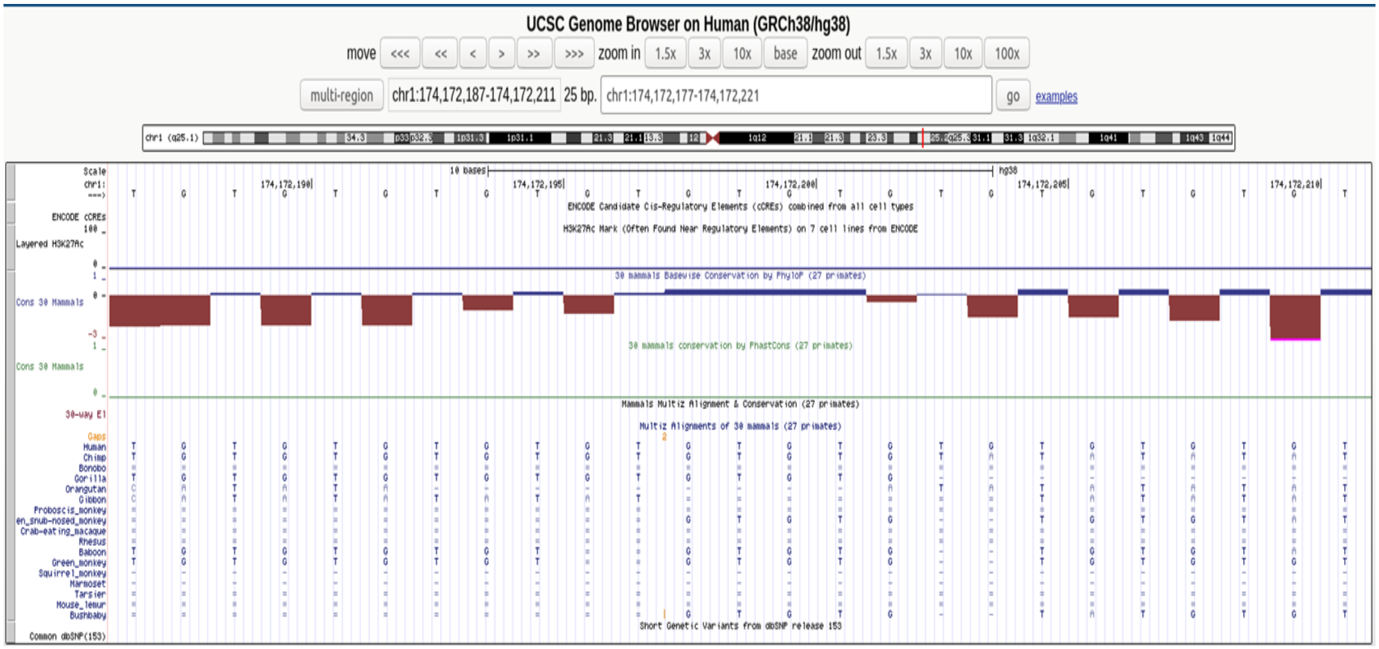

**Case F:** Elongation of CA repeats in intronic region of HSD11B2 gene. Except Orangutan, incomplete CA repeats are observed in all other primates as compared to human, indicating selection and elongation of CA repeats in human HSD11B2 gene.

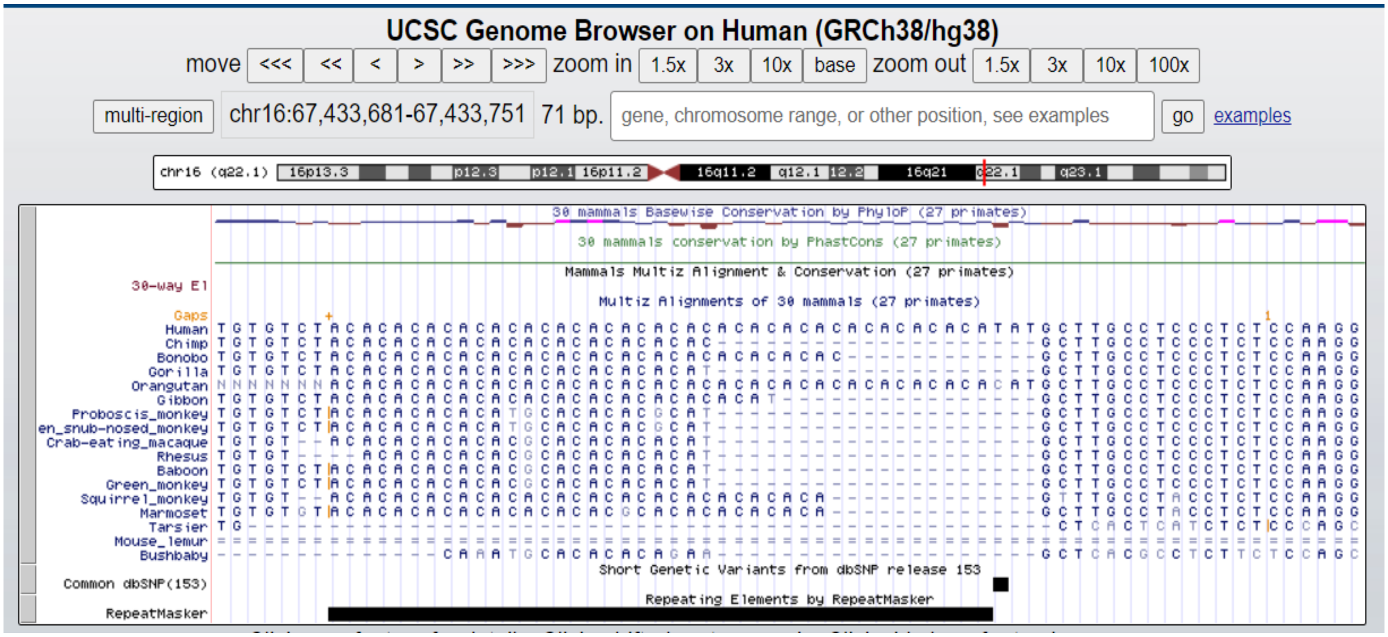
