## Supplementary tables for "Comparative Analysis of TG/CA Repeats in Sixteen Primate Genomes Reveals the Dynamics and Role of TG/CA Repeats in the Human Genome"

Supplementary table S1: Information about primate species used for the analysis

| Common Name | Scientific Name | Abbreviation | Assembly Identifier | Percentage Sequenced |
|---|---|---|---|---|
| Bushbaby | Otolemur garnettii | BB | Otolemur garnettii, build, otoGar3, Mar. 2011 | 93.64 |
| Mouse Lemur | Microcebus murinus | ML | Microcebus murinus, build micMur3, May. 2015 | 95.94 |
| Tarsier | Tarsius syrichta | TS | Tarsius syrichta, build tarSyr2, Sep. 2013 | 98.61 |
| Squirell Monkey | Saimiri boliviensis | SM | Saimiri boliviensis, build saiBol1, Oct. 2011 | 94.96 |
| Marmoset | Callithrix jacchus | MS | Callithrix jacchus, build calJac3, Mar. 2009 | 94.43 |
| Golden snub-nosed monkey | Rhinopithecus roxellana | GS | Rhinopithecus roxellana, build rhiRox1, Oct. 2014 | 98.50 |
| Proboscis Monkey | Nasalis larvatus | PM | Nasalis larvatus, build nasLar1, Nov. 2014 | 79.61 |
| Baboon | Papio anubis | BN | Papio anubis, build papAnu4, Apr. 2017 | 99.24 |
| Rhesus | Macaca mulatta | RH | Macaca mulatta, build rheMac8, Nov. 2015 | 97.09 |
| Crab-eating Macaque | Macaca fascicularis | CM | Macaca fascicularis, build macFas5, Jun. 2013 | 95.15 |
| Green Monkey | Chlorocebus sabaeus | GM | Chlorocebus sabaeus, build ChlSab2, Mar. 2014 | 98.65 |
| Gibbon | Nomascus leucogenys | GB | Nomascus leucogenys, build NomLeu3, Oct. 2012 | 93.06 |
| Orangatun | Pongo pygmaeus abelii | OG | Pongo pygmaeus abelii, build ponAbe3, Jan. 2018 | 99.29 |
| Gorilla | Gorilla gorilla gorilla | GR | Gorilla gorilla gorilla, build gorGor5, Mar. 2016 | 100 |
| Bonobo | Pan paniscus | BO | Pan paniscus, build panPan2, Aug. 2015 | 82.94 |
| Chimpanzee | Pan troglodytes | CP | Pan troglodytes, build panTro6, Jan. 2018 | 98.96 |

Supplementary table S2: Step conversion for other dinucleotide repeats from non-human primate genomes which got converted to TG repeats in human genome

| Repeat | CPZ | BNB | GR | OG | GB | GM | CM | RM | BBN | PM | GSM | MMT | SM | TSR | ML | BB |
|---|---|---|---|---|---|---|---|---|---|---|---|---|---|---|---|---|
| (AA)1 | 89 | 100 | 98 | 270 | 315 | 434 | 611 | 440 | 760 | 470 | 624 | 1057 | 921 | 1812 | 97 | 2204 |
| (AA)2 | 8 | 4 | 11 | 13 | 19 | 29 | 33 | 29 | 36 | 20 | 26 | 39 | 34 | 85 | 6 | 104 |
| (AA)3 | 1 | 0 | 2 | 4 | 2 | 1 | 6 | 10 | 10 | 3 | 3 | 6 | 4 | 17 | 0 | 16 |
| (AA)4 | 0 | 0 | 1 | 0 | 1 | 3 | 0 | 5 | 2 | 0 | 0 | 3 | 1 | 3 | 0 | 1 |
| (AA)5 | 0 | 0 | 0 | 2 | 1 | 1 | 0 | 1 | 0 | 1 | 0 | 2 | 0 | 1 | 0 | 1 |
| (AA)6 | 0 | 0 | 0 | 0 | 0 | 0 | 0 | 0 | 0 | 0 | 0 | 0 | 1 | 2 | 0 | 2 |
| (AA)7 | 0 | 0 | 0 | 0 | 0 | 0 | 0 | 0 | 1 | 0 | 0 | 0 | 0 | 0 | 0 | 0 |
| (AA)8 | 0 | 0 | 0 | 0 | 0 | 0 | 0 | 1 | 0 | 0 | 0 | 0 | 0 | 0 | 0 | 0 |
| (AC)1 | 42 | 53 | 45 | 105 | 83 | 185 | 176 | 184 | 247 | 149 | 177 | 221 | 207 | 357 | 36 | 631 |
| (AC)2 | 2 | 1 | 0 | 2 | 0 | 1 | 2 | 2 | 3 | 0 | 0 | 0 | 0 | 3 | 0 | 9 |
| (AC)3 | 0 | 0 | 0 | 0 | 0 | 1 | 0 | 0 | 0 | 0 | 0 | 0 | 0 | 0 | 0 | 0 |
| (AC)4 | 0 | 0 | 0 | 0 | 0 | 1 | 0 | 0 | 0 | 0 | 0 | 0 | 0 | 0 | 0 | 0 |
| (AG)1 | 1351 | 1336 | 1398 | 2190 | 2332 | 3009 | 3409 | 2852 | 3658 | 2802 | 3471 | 4311 | 3797 | 4245 | 463 | 5676 |
| (AG)10 | 1 | 1 | 4 | 6 | 3 | 16 | 14 | 10 | 23 | 4 | 10 | 21 | 25 | 10 | 2 | 5 |
| (AG)11 | 1 | 1 | 2 | 1 | 3 | 9 | 9 | 5 | 12 | 3 | 13 | 20 | 20 | 12 | 0 | 4 |
| (AG)12 | 2 | 2 | 1 | 1 | 3 | 5 | 3 | 5 | 4 | 4 | 8 | 11 | 15 | 6 | 0 | 2 |
| (AG)13 | 2 | 0 | 0 | 0 | 2 | 3 | 1 | 3 | 3 | 2 | 4 | 10 | 12 | 8 | 0 | 2 |
| (AG)14 | 0 | 0 | 1 | 1 | 0 | 3 | 2 | 1 | 3 | 0 | 5 | 11 | 14 | 11 | 0 | 1 |
| (AG)15 | 0 | 0 | 0 | 0 | 1 | 1 | 1 | 2 | 1 | 1 | 2 | 10 | 6 | 2 | 1 | 1 |
| (AG)16 | 0 | 0 | 0 | 0 | 0 | 0 | 1 | 2 | 1 | 0 | 5 | 8 | 6 | 0 | 0 | 0 |
| (AG)17 | 0 | 0 | 1 | 0 | 0 | 0 | 1 | 0 | 0 | 2 | 2 | 5 | 6 | 3 | 0 | 1 |
| (AG)18 | 0 | 0 | 0 | 0 | 0 | 0 | 1 | 1 | 0 | 0 | 1 | 2 | 1 | 4 | 0 | 0 |
| (AG)19 | 0 | 0 | 0 | 0 | 0 | 1 | 0 | 0 | 0 | 0 | 0 | 0 | 2 | 3 | 0 | 0 |
| (AG)2 | 179 | 190 | 183 | 239 | 219 | 404 | 353 | 340 | 421 | 250 | 358 | 399 | 392 | 376 | 43 | 559 |
| (AG)20 | 0 | 0 | 0 | 0 | 0 | 0 | 0 | 0 | 0 | 0 | 1 | 0 | 0 | 1 | 0 | 0 |
| (AG)21 | 0 | 0 | 0 | 0 | 0 | 0 | 1 | 0 | 0 | 0 | 0 | 1 | 0 | 1 | 0 | 0 |
| (AG)22 | 0 | 0 | 0 | 0 | 0 | 0 | 0 | 0 | 0 | 0 | 0 | 1 | 0 | 0 | 0 | 0 |
| (AG)23 | 0 | 0 | 0 | 0 | 0 | 0 | 1 | 0 | 0 | 0 | 1 | 0 | 0 | 0 | 0 | 0 |
| (AG)25 | 0 | 0 | 0 | 0 | 0 | 0 | 0 | 0 | 0 | 0 | 0 | 1 | 0 | 0 | 0 | 0 |
| (AG)3 | 88 | 76 | 97 | 102 | 119 | 199 | 172 | 177 | 223 | 142 | 189 | 205 | 195 | 141 | 20 | 227 |
| (AG)4 | 65 | 51 | 48 | 86 | 65 | 133 | 112 | 107 | 132 | 94 | 116 | 155 | 137 | 86 | 13 | 131 |
| (AG)5 | 33 | 36 | 40 | 52 | 47 | 87 | 101 | 80 | 112 | 53 | 88 | 122 | 103 | 50 | 8 | 91 |
| (AG)6 | 22 | 23 | 24 | 31 | 44 | 89 | 82 | 69 | 96 | 51 | 80 | 104 | 99 | 61 | 4 | 50 |

| | | | | | | | | | | | | | | | |
|---|---|---|---|---|---|---|---|---|---|---|---|---|---|---|---|
| (AG)7 | 11 | 5 | 11 | 27 | 19 | 47 | 56 | 37 | 60 | 38 | 71 | 54 | 53 | 39 | 2 | 31 |
| (AG)8 | 4 | 7 | 17 | 11 | 8 | 41 | 37 | 20 | 47 | 22 | 38 | 45 | 54 | 32 | 4 | 20 |
| (AG)9 | 8 | 4 | 8 | 13 | 11 | 24 | 24 | 24 | 23 | 9 | 33 | 39 | 36 | 20 | 0 | 12 |
| (AT)1 | 34 | 37 | 47 | 84 | 77 | 178 | 165 | 143 | 289 | 127 | 200 | 259 | 246 | 372 | 32 | 683 |
| (AT)2 | 1 | 1 | 0 | 0 | 0 | 1 | 0 | 3 | 3 | 0 | 0 | 1 | 0 | 4 | 0 | 7 |
| (CA)1 | 732 | 713 | 850 | 1910 | 2545 | 3473 | 4250 | 3284 | 4782 | 3797 | 4464 | 6625 | 5649 | 6695 | 567 | 8189 |
| (CA)10 | 0 | 0 | 0 | 2 | 1 | 4 | 7 | 0 | 8 | 2 | 4 | 6 | 9 | 8 | 0 | 1 |
| (CA)11 | 0 | 0 | 0 | 0 | 1 | 2 | 1 | 0 | 5 | 2 | 5 | 9 | 12 | 9 | 0 | 2 |
| (CA)12 | 0 | 0 | 0 | 0 | 0 | 1 | 2 | 0 | 3 | 0 | 0 | 2 | 6 | 5 | 0 | 3 |
| (CA)13 | 0 | 0 | 0 | 0 | 1 | 0 | 2 | 0 | 5 | 2 | 1 | 5 | 7 | 1 | 0 | 1 |
| (CA)14 | 0 | 0 | 0 | 0 | 0 | 1 | 0 | 0 | 1 | 3 | 0 | 0 | 1 | 0 | 0 | 0 |
| (CA)15 | 0 | 0 | 0 | 0 | 0 | 0 | 1 | 1 | 0 | 0 | 0 | 1 | 2 | 2 | 0 | 0 |
| (CA)16 | 0 | 0 | 0 | 0 | 0 | 0 | 0 | 1 | 0 | 0 | 1 | 2 | 0 | 3 | 0 | 0 |
| (CA)17 | 0 | 0 | 0 | 0 | 0 | 0 | 1 | 0 | 0 | 1 | 0 | 1 | 0 | 5 | 0 | 0 |
| (CA)18 | 0 | 0 | 0 | 0 | 0 | 0 | 0 | 0 | 0 | 0 | 0 | 0 | 0 | 2 | 0 | 0 |
| (CA)19 | 0 | 0 | 0 | 0 | 0 | 0 | 0 | 0 | 0 | 1 | 0 | 0 | 0 | 1 | 0 | 0 |
| (CA)2 | 24 | 27 | 29 | 72 | 181 | 186 | 296 | 185 | 369 | 240 | 262 | 488 | 428 | 572 | 41 | 931 |
| (CA)20 | 0 | 0 | 0 | 0 | 0 | 0 | 0 | 0 | 0 | 0 | 0 | 1 | 0 | 0 | 0 | 0 |
| (CA)21 | 0 | 0 | 0 | 0 | 0 | 0 | 0 | 0 | 0 | 0 | 0 | 0 | 0 | 2 | 0 | 0 |
| (CA)22 | 0 | 0 | 0 | 0 | 0 | 0 | 1 | 0 | 0 | 0 | 0 | 0 | 0 | 0 | 0 | 0 |
| (CA)3 | 7 | 2 | 13 | 13 | 48 | 54 | 85 | 41 | 102 | 66 | 84 | 111 | 117 | 152 | 2 | 182 |
| (CA)4 | 4 | 4 | 3 | 6 | 14 | 25 | 51 | 22 | 52 | 36 | 49 | 59 | 64 | 49 | 0 | 57 |
| (CA)5 | 1 | 1 | 2 | 5 | 12 | 12 | 28 | 10 | 38 | 21 | 22 | 33 | 30 | 41 | 0 | 24 |
| (CA)6 | 0 | 1 | 1 | 2 | 3 | 10 | 33 | 21 | 33 | 17 | 33 | 28 | 23 | 37 | 0 | 15 |
| (CA)7 | 0 | 0 | 0 | 1 | 4 | 10 | 17 | 4 | 22 | 9 | 12 | 17 | 15 | 20 | 0 | 7 |
| (CA)8 | 0 | 0 | 1 | 0 | 1 | 2 | 5 | 4 | 13 | 8 | 9 | 15 | 11 | 12 | 0 | 4 |
| (CA)9 | 0 | 0 | 0 | 0 | 2 | 1 | 5 | 2 | 9 | 4 | 8 | 3 | 9 | 6 | 0 | 0 |
| (CC)1 | 40 | 31 | 58 | 119 | 159 | 295 | 314 | 255 | 413 | 270 | 337 | 640 | 556 | 784 | 80 | 1703 |
| (CC)2 | 0 | 0 | 0 | 3 | 2 | 5 | 3 | 5 | 8 | 6 | 2 | 11 | 6 | 13 | 0 | 55 |
| (CC)3 | 0 | 0 | 0 | 2 | 0 | 1 | 1 | 0 | 2 | 0 | 0 | 1 | 3 | 2 | 0 | 2 |
| (CC)4 | 0 | 0 | 0 | 0 | 0 | 1 | 2 | 0 | 0 | 0 | 0 | 0 | 0 | 0 | 0 | 3 |
| (CC)5 | 0 | 0 | 0 | 0 | 0 | 0 | 0 | 1 | 0 | 0 | 0 | 0 | 0 | 0 | 0 | 0 |
| (CG)1 | 4018 | 3864 | 4380 | 6661 | 7272 | 8297 | 8891 | 8209 | 9669 | 7221 | 8526 | 8399 | 8287 | 4732 | 779 | 5818 |
| (CG)10 | 0 | 0 | 0 | 0 | 0 | 0 | 0 | 0 | 2 | 1 | 0 | 2 | 2 | 1 | 0 | 0 |
| (CG)11 | 0 | 0 | 0 | 0 | 0 | 0 | 1 | 0 | 0 | 0 | 0 | 1 | 0 | 0 | 0 | 0 |
| (CG)12 | 0 | 1 | 0 | 0 | 0 | 1 | 0 | 0 | 0 | 0 | 0 | 1 | 0 | 0 | 0 | 0 |
| (CG)2 | 262 | 247 | 205 | 383 | 438 | 916 | 814 | 731 | 951 | 574 | 705 | 712 | 578 | 288 | 53 | 329 |

| | | | | | | | | | | | | | | | | |
|---|---|---|---|---|---|---|---|---|---|---|---|---|---|---|---|---|
| (CG)3 | 119 | 99 | 68 | 116 | 143 | 344 | 334 | 300 | 346 | 203 | 296 | 239 | 204 | 100 | 16 | 64 |
| (CG)4 | 39 | 46 | 32 | 43 | 60 | 165 | 148 | 156 | 180 | 94 | 147 | 116 | 76 | 43 | 11 | 32 |
| (CG)5 | 21 | 22 | 14 | 12 | 28 | 94 | 64 | 53 | 89 | 49 | 56 | 53 | 44 | 27 | 4 | 16 |
| (CG)6 | 6 | 14 | 1 | 6 | 10 | 38 | 33 | 33 | 29 | 24 | 25 | 19 | 14 | 7 | 1 | 8 |
| (CG)7 | 6 | 2 | 2 | 1 | 3 | 10 | 14 | 9 | 10 | 8 | 13 | 9 | 18 | 3 | 81 | 2 |
| (CG)8 | 2 | 1 | 0 | 0 | 0 | 1 | 4 | 4 | 7 | 1 | 5 | 5 | 1 | 3 | 0 | 1 |
| (CG)9 | 0 | 1 | 0 | 0 | 1 | 3 | 0 | 2 | 3 | 2 | 1 | 2 | 0 | 0 | 0 | 0 |
| (CT)1 | 59 | 52 | 60 | 142 | 217 | 326 | 393 | 343 | 570 | 345 | 413 | 624 | 527 | 989 | 0 | 1672 |
| (CT)2 | 0 | 0 | 0 | 0 | 2 | 1 | 4 | 2 | 6 | 1 | 1 | 5 | 2 | 9 | 0 | 30 |
| (CT)3 | 0 | 0 | 1 | 0 | 0 | 0 | 0 | 0 | 0 | 0 | 0 | 1 | 1 | 2 | 0 | 1 |
| (CT)4 | 0 | 0 | 0 | 0 | 0 | 0 | 0 | 0 | 1 | 0 | 0 | 0 | 0 | 0 | 0 | 0 |
| (GA)1 | 59 | 66 | 78 | 170 | 241 | 361 | 444 | 340 | 690 | 395 | 487 | 615 | 535 | 979 | 54 | 1638 |
| (GA)2 | 4 | 3 | 3 | 2 | 5 | 7 | 10 | 7 | 24 | 8 | 7 | 12 | 9 | 30 | 0 | 67 |
| (GA)3 | 0 | 0 | 0 | 1 | 1 | 0 | 0 | 2 | 4 | 0 | 0 | 2 | 2 | 1 | 0 | 10 |
| (GA)4 | 0 | 0 | 0 | 0 | 0 | 0 | 0 | 0 | 0 | 0 | 0 | 0 | 0 | 0 | 0 | 2 |
| (GA)5 | 0 | 0 | 0 | 0 | 0 | 0 | 0 | 0 | 1 | 0 | 0 | 0 | 0 | 0 | 0 | 0 |
| (GC)1 | 32 | 20 | 23 | 51 | 26 | 106 | 93 | 100 | 148 | 57 | 75 | 108 | 98 | 160 | 16 | 455 |
| (GG)1 | 1276 | 1160 | 1379 | 2168 | 1930 | 3050 | 3152 | 3025 | 3869 | 2352 | 3073 | 3460 | 3354 | 2997 | 313 | 5320 |
| (GG)10 | 0 | 0 | 0 | 0 | 0 | 1 | 0 | 0 | 0 | 0 | 0 | 0 | 0 | 0 | 0 | 0 |
| (GG)11 | 0 | 0 | 0 | 0 | 0 | 0 | 0 | 0 | 0 | 0 | 0 | 0 | 1 | 0 | 0 | 0 |
| (GG)12 | 0 | 0 | 0 | 0 | 0 | 0 | 0 | 0 | 0 | 0 | 1 | 0 | 0 | 0 | 0 | 0 |
| (GG)13 | 0 | 0 | 0 | 0 | 0 | 1 | 0 | 0 | 0 | 0 | 0 | 0 | 0 | 0 | 0 | 0 |
| (GG)2 | 168 | 125 | 130 | 332 | 244 | 544 | 545 | 537 | 695 | 361 | 454 | 520 | 537 | 409 | 52 | 832 |
| (GG)3 | 45 | 21 | 32 | 94 | 70 | 126 | 167 | 154 | 195 | 123 | 151 | 129 | 135 | 77 | 11 | 141 |
| (GG)4 | 15 | 17 | 17 | 27 | 12 | 33 | 34 | 44 | 67 | 38 | 35 | 37 | 33 | 14 | 2 | 44 |
| (GG)5 | 14 | 2 | 10 | 11 | 13 | 13 | 14 | 13 | 42 | 11 | 20 | 12 | 6 | 3 | 0 | 16 |
| (GG)6 | 0 | 0 | 8 | 4 | 2 | 6 | 2 | 9 | 5 | 5 | 3 | 2 | 0 | 0 | 0 | 1 |
| (GG)7 | 1 | 1 | 3 | 2 | 1 | 2 | 0 | 1 | 5 | 0 | 5 | 1 | 0 | 0 | 0 | 0 |
| (GG)8 | 0 | 0 | 4 | 0 | 0 | 1 | 0 | 0 | 0 | 1 | 0 | 1 | 0 | 1 | 1 | 1 |
| (GG)9 | 0 | 0 | 2 | 1 | 1 | 2 | 0 | 0 | 0 | 0 | 3 | 0 | 0 | 0 | 0 | 0 |
| (GT)1 | 68 | 114 | 123 | 155 | 85 | 229 | 187 | 207 | 517 | 153 | 191 | 176 | 205 | 201 | 25 | 615 |
| (GT)2 | 0 | 0 | 0 | 0 | 0 | 0 | 0 | 0 | 3 | 0 | 0 | 0 | 0 | 0 | 0 | 1 |
| (TA)1 | 8138 | 8024 | 8629 | 13476 | 13245 | 16375 | 18043 | 15808 | 19751 | 15646 | 19277 | 17564 | 17104 | 14822 | 1801 | 16293 |
| (TA)10 | 14 | 12 | 13 | 14 | 33 | 2 | 15 | 13 | 12 | 10 | 20 | 19 | 23 | 2 | 0 | 6 |
| (TA)11 | 12 | 11 | 10 | 8 | 23 | 6 | 6 | 3 | 10 | 10 | 11 | 16 | 18 | 7 | 0 | 2 |
| (TA)12 | 5 | 5 | 11 | 6 | 13 | 0 | 3 | 3 | 8 | 7 | 6 | 14 | 7 | 2 | 0 | 1 |

| | | | | | | | | | | | | | | | | |
|---|---|---|---|---|---|---|---|---|---|---|---|---|---|---|---|---|
| (TA)13 | 1 | 3 | 0 | 5 | 12 | 4 | 3 | 0 | 4 | 5 | 8 | 10 | 9 | 4 | 0 | 0 |
| (TA)14 | 4 | 2 | 1 | 3 | 5 | 4 | 1 | 1 | 6 | 1 | 7 | 9 | 6 | 2 | 0 | 0 |
| (TA)15 | 0 | 2 | 1 | 0 | 6 | 0 | 1 | 1 | 1 | 2 | 3 | 3 | 2 | 2 | 0 | 0 |
| (TA)16 | 0 | 0 | 0 | 4 | 1 | 1 | 1 | 2 | 1 | 2 | 0 | 5 | 5 | 2 | 0 | 0 |
| (TA)17 | 0 | 1 | 1 | 0 | 1 | 0 | 1 | 0 | 0 | 0 | 3 | 2 | 1 | 1 | 0 | 0 |
| (TA)18 | 0 | 1 | 1 | 2 | 3 | 0 | 0 | 0 | 1 | 0 | 0 | 2 | 1 | 2 | 194 | 0 |
| (TA)19 | 0 | 1 | 0 | 3 | 1 | 0 | 0 | 0 | 0 | 2 | 2 | 0 | 1 | 0 | 0 | 0 |
| (TA)2 | 977 | 1023 | 1135 | 1537 | 1749 | 1752 | 2422 | 1685 | 2666 | 1939 | 2437 | 2344 | 2170 | 2531 | 0 | 2441 |
| (TA)20 | 0 | 0 | 0 | 1 | 1 | 0 | 1 | 1 | 0 | 0 | 0 | 0 | 1 | 0 | 0 | 0 |
| (TA)21 | 0 | 0 | 0 | 1 | 0 | 0 | 0 | 0 | 0 | 0 | 0 | 0 | 0 | 0 | 0 | 0 |
| (TA)25 | 0 | 1 | 0 | 0 | 0 | 0 | 0 | 0 | 0 | 0 | 0 | 0 | 0 | 0 | 0 | 0 |
| (TA)3 | 453 | 436 | 492 | 731 | 880 | 613 | 1088 | 645 | 1117 | 815 | 1010 | 843 | 806 | 801 | 48 | 677 |
| (TA)4 | 274 | 275 | 307 | 420 | 575 | 340 | 598 | 339 | 608 | 440 | 559 | 423 | 383 | 222 | 16 | 227 |
| (TA)5 | 167 | 152 | 189 | 296 | 325 | 147 | 269 | 166 | 270 | 225 | 262 | 159 | 171 | 77 | 9 | 54 |
| (TA)6 | 107 | 104 | 105 | 152 | 187 | 71 | 132 | 88 | 140 | 116 | 166 | 115 | 114 | 40 | 0 | 24 |
| (TA)7 | 65 | 54 | 89 | 81 | 119 | 36 | 76 | 33 | 76 | 59 | 86 | 56 | 49 | 18 | 2 | 10 |
| (TA)8 | 45 | 41 | 35 | 44 | 82 | 18 | 40 | 29 | 38 | 32 | 47 | 49 | 38 | 14 | 0 | 11 |
| (TA)9 | 23 | 17 | 18 | 27 | 34 | 9 | 23 | 13 | 20 | 18 | 30 | 18 | 27 | 9 | 0 | 5 |
| (TC)1 | 2150 | 2098 | 2331 | 3816 | 3627 | 5124 | 5042 | 4816 | 5535 | 4047 | 5050 | 4991 | 4751 | 3922 | 558 | 5795 |
| (TC)10 | 2 | 3 | 4 | 2 | 10 | 7 | 9 | 11 | 11 | 13 | 10 | 8 | 11 | 6 | 1 | 1 |
| (TC)11 | 2 | 2 | 2 | 10 | 3 | 11 | 4 | 6 | 3 | 3 | 9 | 6 | 8 | 0 | 0 | 0 |
| (TC)12 | 3 | 0 | 0 | 0 | 2 | 4 | 4 | 2 | 3 | 4 | 10 | 9 | 3 | 1 | 0 | 0 |
| (TC)13 | 0 | 1 | 1 | 0 | 0 | 5 | 3 | 4 | 3 | 2 | 9 | 2 | 1 | 3 | 0 | 0 |
| (TC)14 | 0 | 0 | 0 | 0 | 1 | 5 | 2 | 1 | 2 | 1 | 4 | 6 | 2 | 5 | 0 | 1 |
| (TC)15 | 0 | 0 | 0 | 0 | 0 | 4 | 0 | 2 | 1 | 1 | 1 | 5 | 4 | 1 | 0 | 0 |
| (TC)16 | 0 | 0 | 0 | 0 | 0 | 1 | 1 | 0 | 0 | 2 | 1 | 1 | 4 | 0 | 0 | 0 |
| (TC)17 | 1 | 0 | 0 | 0 | 0 | 1 | 0 | 0 | 0 | 1 | 2 | 1 | 0 | 0 | 0 | 0 |
| (TC)18 | 0 | 0 | 0 | 0 | 0 | 0 | 1 | 1 | 0 | 1 | 2 | 0 | 0 | 0 | 0 | 0 |
| (TC)19 | 0 | 0 | 0 | 0 | 0 | 1 | 0 | 0 | 0 | 1 | 1 | 1 | 0 | 0 | 0 | 0 |
| (TC)2 | 138 | 129 | 144 | 195 | 188 | 330 | 294 | 314 | 1 | 225 | 311 | 273 | 250 | 273 | 22 | 470 |
| (TC)20 | 0 | 0 | 0 | 0 | 0 | 0 | 0 | 0 | 361 | 1 | 0 | 0 | 0 | 0 | 0 | 0 |
| (TC)21 | 0 | 0 | 0 | 0 | 0 | 0 | 1 | 0 | 0 | 0 | 1 | 0 | 0 | 0 | 0 | 0 |
| (TC)22 | 0 | 0 | 0 | 0 | 0 | 0 | 0 | 0 | 0 | 1 | 0 | 1 | 1 | 0 | 0 | 0 |

| | | | | | | | | | | | | | | | | |
|---|---|---|---|---|---|---|---|---|---|---|---|---|---|---|---|---|
| (TC)23 | 0 | 0 | 0 | 0 | 0 | 0 | 0 | 0 | 0 | 0 | 1 | 0 | 0 | 0 | 0 | 0 |
| (TC)3 | 82 | 53 | 56 | 94 | 71 | 83 | 100 | 103 | 1 | 77 | 121 | 96 | 90 | 80 | 4 | 66 |
| (TC)4 | 44 | 30 | 36 | 44 | 50 | 64 | 62 | 56 | 130 | 42 | 70 | 53 | 54 | 47 | 0 | 28 |
| (TC)5 | 19 | 22 | 23 | 30 | 44 | 48 | 52 | 38 | 79 | 34 | 55 | 34 | 33 | 22 | 1 | 20 |
| (TC)6 | 21 | 10 | 18 | 33 | 31 | 34 | 42 | 36 | 48 | 30 | 56 | 23 | 33 | 27 | 3 | 14 |
| (TC)7 | 7 | 11 | 11 | 13 | 17 | 31 | 25 | 29 | 41 | 17 | 43 | 31 | 18 | 14 | 2 | 4 |
| (TC)8 | 6 | 9 | 4 | 5 | 18 | 21 | 22 | 16 | 20 | 15 | 22 | 21 | 18 | 10 | 1 | 4 |
| (TC)9 | 2 | 5 | 6 | 6 | 12 | 18 | 12 | 11 | 24 | 10 | 24 | 12 | 10 | 5 | 0 | 0 |
| (TT)1 | 2794 | 2606 | 3111 | 5328 | 4972 | 6608 | 6837 | 6194 | 3 | 5511 | 6863 | 7222 | 6679 | 6173 | 878 | 8164 |
| (TT)10 | 3 | 0 | 5 | 9 | 11 | 15 | 15 | 18 | 8040 | 7 | 13 | 3 | 7 | 0 | 1 | 2 |
| (TT)11 | 1 | 0 | 5 | 11 | 4 | 7 | 9 | 16 | 16 | 4 | 12 | 3 | 6 | 0 | 0 | 1 |
| (TT)12 | 1 | 0 | 3 | 5 | 5 | 6 | 7 | 3 | 8 | 4 | 6 | 1 | 7 | 0 | 0 | 0 |
| (TT)13 | 1 | 0 | 2 | 1 | 6 | 5 | 4 | 4 | 7 | 2 | 4 | 0 | 3 | 0 | 0 | 0 |
| (TT)14 | 1 | 0 | 0 | 0 | 2 | 2 | 3 | 1 | 2 | 1 | 6 | 1 | 2 | 0 | 0 | 0 |
| (TT)15 | 0 | 1 | 3 | 1 | 0 | 0 | 0 | 0 | 2 | 0 | 2 | 1 | 0 | 0 | 0 | 0 |
| (TT)16 | 0 | 0 | 0 | 1 | 0 | 0 | 1 | 1 | 1 | 0 | 2 | 0 | 1 | 0 | 0 | 0 |
| (TT)17 | 0 | 0 | 0 | 0 | 1 | 0 | 1 | 0 | 0 | 1 | 1 | 0 | 0 | 0 | 0 | 0 |
| (TT)18 | 0 | 0 | 0 | 0 | 0 | 0 | 1 | 1 | 0 | 0 | 0 | 0 | 0 | 0 | 0 | 0 |
| (TT)19 | 0 | 0 | 0 | 0 | 0 | 0 | 1 | 1 | 0 | 0 | 0 | 0 | 0 | 0 | 0 | 0 |
| (TT)2 | 225 | 211 | 271 | 587 | 588 | 724 | 871 | 714 | 2 | 582 | 848 | 953 | 918 | 785 | 99 | 1101 |
| (TT)21 | 0 | 0 | 0 | 0 | 0 | 1 | 0 | 0 | 0 | 0 | 0 | 0 | 0 | 0 | 0 | 0 |
| (TT)3 | 104 | 73 | 109 | 283 | 284 | 316 | 385 | 296 | 1119 | 246 | 309 | 305 | 326 | 173 | 33 | 233 |
| (TT)4 | 50 | 33 | 68 | 166 | 143 | 135 | 205 | 148 | 449 | 128 | 140 | 130 | 152 | 64 | 11 | 66 |
| (TT)5 | 38 | 21 | 48 | 94 | 113 | 93 | 134 | 115 | 215 | 66 | 125 | 84 | 107 | 35 | 6 | 44 |
| (TT)6 | 20 | 8 | 45 | 70 | 78 | 79 | 108 | 88 | 171 | 57 | 107 | 58 | 64 | 9 | 5 | 36 |
| (TT)7 | 15 | 1 | 29 | 47 | 44 | 52 | 56 | 56 | 127 | 27 | 58 | 33 | 38 | 6 | 1 | 13 |
| (TT)8 | 10 | 2 | 12 | 33 | 31 | 33 | 43 | 31 | 71 | 25 | 41 | 12 | 24 | 2 | 1 | 8 |
| (TT)9 | 5 | 2 | 5 | 17 | 15 | 16 | 30 | 23 | 61 | 15 | 19 | 8 | 14 | 1 | 0 | 1 |

Supplementary table S3: Step conversion for other dinucleotide repeats from non-human primate genomes which got converted to CA repeats in human genome

| Repeat | CPZ | BNB | GR | OG | GB | GM | CM | RM | BBN | PM | GSM | MMT | SM | TSR | ML | BB |
|---|---|---|---|---|---|---|---|---|---|---|---|---|---|---|---|---|
| (AA)1 | 2810 | 2652 | 3170 | 5267 | 4811 | 6371 | 7020 | 6303 | 8199 | 5337 | 6928 | 7208 | 6752 | 6159 | 852 | 7875 |
| (AA)10 | 4 | 1 | 4 | 25 | 12 | 17 | 17 | 20 | 21 | 6 | 26 | 6 | 6 | 1 | 106 | 2 |
| (AA)11 | 0 | 0 | 3 | 6 | 7 | 10 | 11 | 5 | 11 | 5 | 15 | 5 | 5 | 0 | 0 | 2 |
| (AA)12 | 0 | 0 | 2 | 1 | 1 | 3 | 8 | 7 | 10 | 5 | 12 | 1 | 1 | 0 | 0 | 0 |
| (AA)13 | 0 | 0 | 0 | 0 | 1 | 0 | 3 | 3 | 3 | 0 | 4 | 0 | 1 | 0 | 0 | 0 |
| (AA)14 | 1 | 0 | 2 | 2 | 2 | 1 | 2 | 4 | 3 | 2 | 5 | 3 | 1 | 0 | 0 | 0 |
| (AA)15 | 0 | 0 | 1 | 0 | 0 | 3 | 2 | 0 | 2 | 1 | 2 | 0 | 0 | 0 | 0 | 0 |
| (AA)16 | 0 | 0 | 0 | 1 | 0 | 1 | 3 | 1 | 3 | 2 | 1 | 0 | 0 | 0 | 0 | 0 |
| (AA)17 | 0 | 0 | 0 | 0 | 2 | 2 | 0 | 0 | 1 | 0 | 2 | 0 | 1 | 0 | 0 | 0 |
| (AA)18 | 0 | 0 | 1 | 1 | 0 | 0 | 0 | 0 | 1 | 0 | 1 | 0 | 1 | 0 | 0 | 0 |
| (AA)19 | 0 | 0 | 0 | 1 | 0 | 1 | 1 | 0 | 0 | 1 | 0 | 0 | 1 | 0 | 0 | 0 |
| (AA)2 | 259 | 164 | 282 | 520 | 570 | 771 | 946 | 791 | 1164 | 698 | 931 | 968 | 888 | 790 | 0 | 1106 |
| (AA)20 | 0 | 0 | 0 | 0 | 0 | 0 | 0 | 0 | 0 | 2 | 0 | 0 | 0 | 0 | 0 | 0 |
| (AA)22 | 0 | 0 | 0 | 0 | 0 | 1 | 0 | 0 | 0 | 0 | 0 | 0 | 0 | 0 | 0 | 0 |
| (AA)3 | 77 | 77 | 109 | 256 | 250 | 288 | 389 | 296 | 445 | 247 | 329 | 303 | 310 | 186 | 30 | 259 |
| (AA)4 | 61 | 27 | 60 | 161 | 148 | 133 | 196 | 141 | 209 | 142 | 152 | 131 | 122 | 50 | 9 | 92 |
| (AA)5 | 33 | 20 | 43 | 89 | 100 | 111 | 117 | 99 | 156 | 75 | 107 | 85 | 72 | 26 | 5 | 52 |
| (AA)6 | 22 | 12 | 32 | 85 | 75 | 99 | 105 | 95 | 147 | 80 | 100 | 66 | 76 | 12 | 1 | 27 |
| (AA)7 | 14 | 1 | 21 | 61 | 48 | 46 | 56 | 53 | 75 | 40 | 69 | 33 | 45 | 8 | 0 | 17 |
| (AA)8 | 9 | 0 | 12 | 34 | 34 | 27 | 45 | 32 | 37 | 22 | 40 | 20 | 28 | 4 | 1 | 9 |
| (AA)9 | 7 | 0 | 9 | 21 | 21 | 15 | 32 | 37 | 38 | 9 | 26 | 11 | 15 | 3 | 0 | 3 |
| (AC)1 | 71 | 124 | 117 | 173 | 99 | 198 | 192 | 197 | 508 | 174 | 179 | 184 | 198 | 237 | 31 | 607 |
| (AC)2 | 0 | 0 | 0 | 0 | 0 | 0 | 0 | 0 | 0 | 0 | 0 | 0 | 0 | 0 | 0 | 0 |
| (AG)1 | 36 | 47 | 68 | 149 | 206 | 349 | 349 | 321 | 8 | 353 | 408 | 599 | 563 | 1009 | 88 | 1704 |
| (AG)2 | 0 | 0 | 3 | 2 | 4 | 1 | 2 | 1 | 600 | 5 | 3 | 5 | 2 | 13 | 0 | 22 |
| (AG)3 | 0 | 0 | 1 | 0 | 0 | 0 | 1 | 1 | 12 | 0 | 0 | 0 | 0 | 1 | 0 | 3 |
| (AG)5 | 0 | 0 | 0 | 0 | 0 | 0 | 0 | 0 | 3 | 0 | 0 | 0 | 0 | 0 | 0 | 0 |
| (AT)1 | 38 | 42 | 52 | 95 | 77 | 185 | 202 | 171 | 1 | 148 | 212 | 289 | 255 | 456 | 20 | 693 |
| (AT)2 | 1 | 2 | 3 | 2 | 0 | 1 | 1 | 1 | 316 | 0 | 0 | 1 | 1 | 1 | 0 | 8 |
| (CC)1 | 1292 | 1109 | 1388 | 2118 | 1959 | 3026 | 3079 | 2950 | 6 | 2378 | 2992 | 3596 | 3371 | 2999 | 376 | 5200 |
| (CC)11 | 0 | 0 | 0 | 0 | 0 | 0 | 0 | 0 | 0 | 0 | 1 | 0 | 0 | 0 | 0 | 0 |
| (CC)12 | 0 | 0 | 0 | 0 | 0 | 1 | 0 | 0 | 0 | 0 | 0 | 0 | 0 | 0 | 0 | 0 |
| (CC)15 | 0 | 0 | 0 | 0 | 0 | 1 | 0 | 0 | 0 | 0 | 0 | 0 | 0 | 0 | 0 | 0 |
| (CC)2 | 139 | 116 | 140 | 332 | 203 | 506 | 533 | 485 | 3876 | 316 | 418 | 491 | 530 | 391 | 29 | 807 |
| (CC)3 | 55 | 26 | 39 | 97 | 49 | 124 | 164 | 165 | 689 | 128 | 124 | 124 | 148 | 80 | 8 | 136 |
| (CC)4 | 17 | 9 | 16 | 25 | 24 | 46 | 45 | 56 | 189 | 33 | 41 | 30 | 30 | 18 | 0 | 33 |

| | | | | | | | | | | | | | | | | |
|---|---|---|---|---|---|---|---|---|---|---|---|---|---|---|---|---|
| (CC)5 | 9 | 4 | 9 | 9 | 15 | 13 | 12 | 18 | 66 | 5 | 21 | 17 | 7 | 6 | 0 | 21 |
| (CC)6 | 3 | 1 | 5 | 1 | 4 | 8 | 0 | 1 | 33 | 4 | 14 | 2 | 0 | 1 | 0 | 1 |
| (CC)7 | 3 | 0 | 1 | 4 | 0 | 5 | 1 | 2 | 3 | 2 | 2 | 0 | 0 | 2 | 0 | 0 |
| (CC)8 | 1 | 0 | 1 | 1 | 1 | 3 | 1 | 0 | 0 | 1 | 3 | 1 | 0 | 0 | 0 | 0 |
| (CC)9 | 0 | 0 | 1 | 0 | 0 | 0 | 1 | 0 | 0 | 0 | 1 | 0 | 0 | 0 | 0 | 0 |
| (CG)1 | 4053 | 3856 | 4290 | 6528 | 7259 | 8270 | 8727 | 8000 | 2 | 7224 | 8638 | 8326 | 8101 | 4547 | 813 | 5993 |
| (CG)10 | 0 | 0 | 0 | 0 | 0 | 0 | 1 | 1 | 0 | 1 | 2 | 1 | 0 | 0 | 0 | 0 |
| (CG)11 | 0 | 0 | 0 | 0 | 1 | 0 | 0 | 1 | 0 | 0 | 1 | 0 | 0 | 0 | 0 | 0 |
| (CG)2 | 288 | 246 | 228 | 396 | 468 | 802 | 811 | 749 | 9435 | 600 | 679 | 697 | 573 | 308 | 68 | 337 |
| (CG)3 | 99 | 84 | 65 | 102 | 153 | 337 | 284 | 262 | 953 | 236 | 265 | 232 | 201 | 95 | 21 | 69 |
| (CG)4 | 28 | 36 | 33 | 37 | 58 | 191 | 142 | 130 | 349 | 100 | 158 | 93 | 78 | 39 | 8 | 31 |
| (CG)5 | 19 | 22 | 8 | 24 | 19 | 104 | 65 | 66 | 196 | 43 | 55 | 45 | 33 | 18 | 5 | 12 |
| (CG)6 | 7 | 10 | 5 | 8 | 6 | 30 | 28 | 23 | 71 | 14 | 25 | 20 | 17 | 10 | 2 | 2 |
| (CG)7 | 2 | 1 | 2 | 4 | 2 | 14 | 14 | 11 | 29 | 11 | 8 | 14 | 8 | 4 | 0 | 1 |
| (CG)8 | 0 | 1 | 0 | 0 | 4 | 12 | 4 | 3 | 18 | 1 | 4 | 5 | 2 | 2 | 0 | 0 |
| (CG)9 | 0 | 0 | 0 | 0 | 1 | 0 | 3 | 1 | 5 | 0 | 3 | 1 | 1 | 0 | 0 | 0 |
| (CT)1 | 1323 | 1328 | 1414 | 2154 | 2390 | 3078 | 3438 | 2897 | 1 | 2770 | 3436 | 4284 | 3649 | 4287 | 481 | 5572 |
| (CT)10 | 4 | 2 | 5 | 3 | 8 | 10 | 16 | 14 | 3852 | 6 | 19 | 31 | 27 | 14 | 2 | 7 |
| (CT)11 | 2 | 0 | 4 | 1 | 4 | 13 | 10 | 6 | 25 | 5 | 7 | 22 | 21 | 3 | 0 | 6 |
| (CT)12 | 0 | 0 | 2 | 0 | 3 | 5 | 5 | 4 | 6 | 2 | 7 | 20 | 16 | 9 | 0 | 2 |
| (CT)13 | 0 | 0 | 0 | 1 | 0 | 2 | 3 | 2 | 11 | 3 | 2 | 11 | 15 | 7 | 0 | 4 |
| (CT)14 | 0 | 0 | 1 | 0 | 1 | 1 | 0 | 1 | 8 | 3 | 5 | 8 | 5 | 9 | 0 | 1 |
| (CT)15 | 0 | 0 | 0 | 0 | 0 | 2 | 1 | 0 | 0 | 4 | 0 | 9 | 7 | 4 | 0 | 0 |
| (CT)16 | 0 | 0 | 0 | 0 | 1 | 0 | 2 | 0 | 0 | 1 | 4 | 6 | 3 | 3 | 0 | 0 |
| (CT)17 | 0 | 0 | 0 | 0 | 0 | 0 | 2 | 1 | 0 | 1 | 1 | 7 | 6 | 5 | 0 | 0 |
| (CT)18 | 0 | 0 | 0 | 0 | 0 | 0 | 1 | 0 | 0 | 0 | 1 | 5 | 0 | 1 | 0 | 0 |
| (CT)19 | 0 | 0 | 0 | 0 | 0 | 1 | 1 | 1 | 0 | 0 | 1 | 2 | 3 | 0 | 0 | 0 |
| (CT)2 | 166 | 158 | 185 | 218 | 201 | 344 | 352 | 316 | 2 | 266 | 328 | 408 | 388 | 371 | 13 | 588 |
| (CT)20 | 0 | 0 | 0 | 0 | 0 | 0 | 0 | 1 | 421 | 0 | 0 | 1 | 1 | 1 | 0 | 0 |
| (CT)22 | 0 | 0 | 0 | 0 | 0 | 0 | 0 | 0 | 0 | 0 | 1 | 0 | 1 | 0 | 0 | 0 |
| (CT)23 | 0 | 0 | 0 | 0 | 0 | 0 | 0 | 0 | 0 | 0 | 0 | 1 | 0 | 0 | 0 | 0 |
| (CT)25 | 0 | 0 | 0 | 0 | 0 | 0 | 0 | 0 | 0 | 0 | 0 | 1 | 0 | 0 | 0 | 0 |
| (CT)3 | 93 | 82 | 89 | 105 | 111 | 168 | 163 | 162 | 1 | 116 | 164 | 184 | 189 | 158 | 4 | 219 |
| (CT)4 | 48 | 60 | 48 | 82 | 62 | 115 | 125 | 90 | 200 | 76 | 123 | 138 | 138 | 76 | 1 | 133 |
| (CT)5 | 25 | 21 | 30 | 52 | 45 | 88 | 82 | 71 | 133 | 65 | 93 | 120 | 116 | 53 | 8 | 73 |
| (CT)6 | 20 | 21 | 30 | 34 | 41 | 75 | 79 | 79 | 90 | 38 | 82 | 117 | 134 | 55 | 1 | 49 |
| (CT)7 | 12 | 7 | 17 | 14 | 26 | 36 | 44 | 46 | 89 | 32 | 50 | 70 | 75 | 38 | 2 | 36 |
| (CT)8 | 7 | 8 | 9 | 13 | 16 | 55 | 32 | 30 | 49 | 26 | 27 | 74 | 71 | 39 | 572 | 21 |
| (CT)9 | 3 | 4 | 8 | 10 | 14 | 21 | 27 | 24 | 39 | 18 | 31 | 46 | 47 | 20 | 1 | 6 |
| (GA)1 | 2162 | 2172 | 2419 | 3741 | 3653 | 5073 | 4950 | 4751 | 29 | 3979 | 5022 | 5048 | 4943 | 3983 | 28 | 5485 |
| (GA)10 | 3 | 3 | 6 | 8 | 7 | 5 | 16 | 15 | 5536 | 8 | 17 | 10 | 7 | 5 | 0 | 1 |

| | | | | | | | | | | | | | | | | |
|---|---|---|---|---|---|---|---|---|---|---|---|---|---|---|---|---|
| (GA)11 | 3 | 3 | 3 | 3 | 4 | 10 | 7 | 11 | 4 | 5 | 8 | 6 | 6 | 1 | 11 | 0 |
| (GA)1 | 0 | 1 | 1 | 2 | 3 | 8 | 6 | 5 | 6 | 1 | 5 | 8 | 4 | 2 | 0 | 0 |
| (GA)13 | 2 | 1 | 0 | 0 | 4 | 2 | 2 | 2 | 3 | 2 | 11 | 5 | 2 | 1 | 0 | 0 |
| (GA)14 | 1 | 1 | 0 | 0 | 0 | 2 | 1 | 1 | 0 | 3 | 2 | 2 | 1 | 1 | 0 | 0 |
| (GA)15 | 0 | 0 | 0 | 0 | 0 | 1 | 3 | 4 | 4 | 0 | 2 | 3 | 1 | 2 | 0 | 0 |
| (GA)16 | 0 | 0 | 0 | 0 | 0 | 0 | 2 | 2 | 0 | 1 | 3 | 1 | 1 | 1 | 0 | 0 |
| (GA)17 | 0 | 0 | 0 | 0 | 0 | 1 | 0 | 0 | 0 | 0 | 0 | 2 | 1 | 0 | 0 | 0 |
| (GA)18 | 0 | 0 | 0 | 0 | 0 | 0 | 0 | 0 | 3 | 0 | 0 | 0 | 0 | 1 | 0 | 0 |
| (GA)19 | 0 | 0 | 0 | 0 | 0 | 0 | 1 | 0 | 0 | 0 | 1 | 1 | 0 | 0 | 0 | 0 |
| (GA)2 | 137 | 134 | 123 | 203 | 199 | 364 | 297 | 305 | 1 | 230 | 305 | 283 | 296 | 261 | 4 | 388 |
| (GA)20 | 0 | 0 | 0 | 0 | 0 | 0 | 0 | 0 | 0 | 0 | 0 | 0 | 1 | 0 | 0 | 0 |
| (GA)21 | 0 | 0 | 0 | 0 | 0 | 0 | 0 | 0 | 0 | 0 | 2 | 1 | 0 | 0 | 0 | 0 |
| (GA)22 | 0 | 0 | 0 | 0 | 0 | 0 | 0 | 0 | 0 | 0 | 0 | 1 | 0 | 0 | 0 | 0 |
| (GA)23 | 0 | 0 | 0 | 0 | 0 | 0 | 0 | 0 | 0 | 1 | 0 | 0 | 0 | 0 | 0 | 0 |
| (GA)3 | 58 | 68 | 58 | 61 | 80 | 127 | 104 | 114 | 406 | 117 | 112 | 86 | 79 | 69 | 1 | 86 |
| (GA)4 | 40 | 28 | 40 | 57 | 34 | 66 | 71 | 66 | 122 | 45 | 74 | 53 | 45 | 44 | 24 | 26 |
| (GA)5 | 29 | 18 | 21 | 35 | 40 | 51 | 42 | 41 | 97 | 41 | 48 | 33 | 36 | 28 | 83 | 12 |
| (GA)6 | 18 | 19 | 14 | 18 | 24 | 40 | 35 | 46 | 57 | 36 | 49 | 32 | 32 | 19 | 0 | 7 |
| (GA)7 | 8 | 12 | 10 | 23 | 22 | 31 | 23 | 15 | 55 | 22 | 35 | 30 | 19 | 16 | 0 | 16 |
| (GA)8 | 7 | 7 | 5 | 15 | 11 | 29 | 20 | 23 | 27 | 16 | 19 | 19 | 21 | 11 | 0 | 2 |
| (GA)9 | 8 | 3 | 2 | 3 | 11 | 20 | 14 | 9 | 26 | 10 | 31 | 15 | 16 | 3 | 2 | 3 |
| (GC)1 | 24 | 20 | 34 | 40 | 28 | 104 | 85 | 95 | 16 | 42 | 82 | 121 | 98 | 168 | 28 | 511 |
| (GC)2 | 0 | 0 | 0 | 1 | 0 | 0 | 0 | 0 | 160 | 0 | 0 | 0 | 0 | 0 | 0 | 0 |
| (GG)1 | 40 | 44 | 56 | 150 | 170 | 409 | 372 | 315 | 1 | 318 | 382 | 622 | 531 | 881 | 1939 | 1686 |
| (GG)2 | 0 | 1 | 0 | 1 | 3 | 9 | 3 | 4 | 469 | 1 | 3 | 11 | 7 | 18 | 1 | 48 |
| (GG)3 | 0 | 0 | 0 | 0 | 0 | 2 | 1 | 4 | 14 | 0 | 1 | 1 | 0 | 1 | 0 | 3 |
| (GG)4 | 0 | 0 | 0 | 1 | 0 | 0 | 0 | 0 | 0 | 0 | 0 | 0 | 0 | 0 | 0 | 1 |
| (GG)7 | 0 | 0 | 0 | 0 | 0 | 0 | 0 | 1 | 0 | 0 | 0 | 0 | 0 | 0 | 0 | 0 |
| (GT)1 | 48 | 48 | 51 | 110 | 86 | 180 | 140 | 176 | 4 | 145 | 196 | 208 | 195 | 338 | 1 | 584 |
| (GT)2 | 2 | 0 | 0 | 2 | 0 | 5 | 1 | 5 | 245 | 0 | 0 | 0 | 0 | 5 | 0 | 5 |
| (GT)3 | 0 | 0 | 0 | 0 | 0 | 1 | 0 | 0 | 0 | 0 | 0 | 0 | 0 | 0 | 0 | 0 |
| (TA)1 | 7873 | 7827 | 8443 | 13403 | 13167 | 16400 | 18014 | 15715 | 1 | 15425 | 19021 | 17542 | 16918 | 15249 | 220 | 16475 |
| (TA)10 | 15 | 11 | 19 | 13 | 26 | 9 | 5 | 5 | 19716 | 10 | 22 | 19 | 16 | 3 | 0 | 1 |
| (TA)11 | 9 | 8 | 6 | 12 | 19 | 5 | 14 | 4 | 12 | 6 | 11 | 13 | 12 | 3 | 0 | 2 |
| (TA)12 | 6 | 5 | 5 | 5 | 16 | 2 | 3 | 1 | 12 | 6 | 8 | 3 | 12 | 1 | 0 | 0 |
| (TA)13 | 2 | 1 | 1 | 3 | 13 | 2 | 5 | 4 | 1 | 1 | 7 | 11 | 6 | 4 | 59 | 0 |
| (TA)14 | 0 | 0 | 4 | 2 | 8 | 2 | 1 | 2 | 6 | 4 | 1 | 5 | 2 | 1 | 11 | 0 |
| (TA)15 | 2 | 1 | 1 | 3 | 10 | 1 | 5 | 2 | 2 | 2 | 3 | 1 | 3 | 1 | 0 | 0 |
| (TA)16 | 1 | 1 | 1 | 0 | 4 | 0 | 3 | 1 | 1 | 0 | 1 | 0 | 1 | 2 | 0 | 0 |

| | | | | | | | | | | | | | | | | |
|---|---|---|---|---|---|---|---|---|---|---|---|---|---|---|---|---|
| (TA)17 | 1 | 0 | 2 | 1 | 1 | 0 | 4 | 0 | 6 | 0 | 1 | 1 | 2 | 1 | 0 | 0 |
| (TA)18 | 0 | 0 | 2 | 1 | 2 | 0 | 0 | 1 | 1 | 1 | 0 | 3 | 0 | 0 | 0 | 0 |
| (TA)19 | 0 | 0 | 0 | 0 | 2 | 0 | 0 | 0 | 3 | 1 | 1 | 0 | 1 | 0 | 0 | 0 |
| (TA)2 | 968 | 978 | 1006 | 1488 | 1773 | 1722 | 2553 | 1743 | 1 | 2020 | 2421 | 2363 | 2182 | 2569 | 4 | 2378 |
| (TA)20 | 0 | 0 | 0 | 0 | 1 | 0 | 1 | 0 | 0 | 1 | 0 | 0 | 0 | 0 | 0 | 0 |
| (TA)21 | 0 | 0 | 1 | 0 | 0 | 0 | 0 | 0 | 0 | 0 | 0 | 2 | 0 | 0 | 0 | 0 |
| (TA)22 | 0 | 0 | 0 | 0 | 0 | 0 | 1 | 0 | 2699 | 0 | 0 | 0 | 0 | 1 | 0 | 0 |
| (TA)3 | 515 | 437 | 492 | 669 | 871 | 601 | 1123 | 632 | 1 | 836 | 1061 | 812 | 725 | 852 | 3 | 602 |
| (TA)4 | 258 | 243 | 323 | 407 | 510 | 300 | 597 | 339 | 1141 | 444 | 576 | 384 | 366 | 306 | 4 | 179 |
| (TA)5 | 163 | 139 | 184 | 254 | 285 | 169 | 298 | 157 | 635 | 189 | 241 | 179 | 176 | 87 | 1 | 56 |
| (TA)6 | 81 | 86 | 122 | 129 | 197 | 74 | 140 | 83 | 307 | 122 | 163 | 133 | 120 | 53 | 77 | 22 |
| (TA)7 | 78 | 65 | 64 | 92 | 93 | 33 | 74 | 46 | 136 | 48 | 69 | 64 | 70 | 25 | 1 | 17 |
| (TA)8 | 30 | 22 | 47 | 46 | 61 | 18 | 29 | 28 | 65 | 38 | 49 | 36 | 33 | 12 | 597 | 5 |
| (TA)9 | 20 | 17 | 25 | 32 | 44 | 8 | 24 | 13 | 36 | 16 | 22 | 35 | 26 | 7 | 0 | 3 |
| (TC)1 | 65 | 55 | 88 | 174 | 219 | 304 | 404 | 284 | 17 | 362 | 470 | 571 | 549 | 946 | 28 | 1690 |
| (TC)2 | 2 | 3 | 4 | 4 | 2 | 4 | 7 | 4 | 674 | 7 | 10 | 16 | 10 | 27 | 3 | 61 |
| (TC)3 | 0 | 1 | 0 | 0 | 0 | 2 | 1 | 1 | 23 | 0 | 2 | 0 | 0 | 0 | 0 | 3 |
| (TC)4 | 0 | 0 | 0 | 0 | 0 | 0 | 0 | 0 | 2 | 0 | 0 | 0 | 0 | 0 | 0 | 2 |
| (TG)1 | 694 | 654 | 829 | 1856 | 2444 | 3470 | 4288 | 3262 | 2 | 3828 | 4531 | 6701 | 5784 | 6642 | 1 | 8254 |
| (TG)10 | 1 | 1 | 0 | 2 | 1 | 0 | 4 | 3 | 4804 | 3 | 4 | 4 | 8 | 7 | 0 | 4 |
| (TG)11 | 0 | 0 | 0 | 1 | 0 | 5 | 1 | 0 | 6 | 3 | 2 | 2 | 8 | 6 | 0 | 2 |
| (TG)12 | 0 | 0 | 0 | 0 | 0 | 0 | 6 | 1 | 5 | 1 | 3 | 0 | 0 | 6 | 0 | 1 |
| (TG)13 | 0 | 0 | 0 | 0 | 1 | 0 | 0 | 2 | 1 | 2 | 4 | 1 | 5 | 6 | 0 | 0 |
| (TG)14 | 0 | 0 | 0 | 0 | 0 | 0 | 0 | 0 | 2 | 0 | 0 | 1 | 2 | 7 | 0 | 0 |
| (TG)15 | 0 | 0 | 0 | 0 | 0 | 2 | 1 | 0 | 1 | 1 | 0 | 0 | 1 | 0 | 0 | 0 |
| (TG)16 | 0 | 0 | 0 | 0 | 0 | 0 | 0 | 0 | 0 | 1 | 0 | 1 | 2 | 2 | 0 | 1 |
| (TG)17 | 0 | 0 | 0 | 0 | 0 | 0 | 0 | 0 | 0 | 0 | 0 | 0 | 1 | 1 | 0 | 0 |
| (TG)18 | 0 | 0 | 0 | 0 | 0 | 0 | 0 | 0 | 0 | 0 | 0 | 0 | 0 | 1 | 0 | 0 |
| (TG)19 | 0 | 0 | 0 | 0 | 0 | 0 | 0 | 0 | 0 | 1 | 0 | 0 | 0 | 1 | 0 | 0 |
| (TG)2 | 24 | 14 | 32 | 90 | 162 | 229 | 340 | 193 | 1 | 271 | 293 | 495 | 452 | 633 | 108 | 883 |
| (TG)20 | 0 | 0 | 0 | 0 | 0 | 0 | 0 | 0 | 0 | 0 | 0 | 0 | 1 | 1 | 0 | 0 |
| (TG)21 | 0 | 0 | 0 | 0 | 0 | 0 | 0 | 0 | 0 | 0 | 1 | 0 | 0 | 0 | 0 | 0 |
| (TG)3 | 6 | 6 | 6 | 12 | 43 | 46 | 101 | 43 | 393 | 57 | 93 | 116 | 108 | 130 | 4 | 159 |
| (TG)4 | 3 | 1 | 3 | 6 | 13 | 33 | 56 | 30 | 103 | 34 | 61 | 73 | 61 | 52 | 0 | 59 |
| (TG)5 | 2 | 1 | 3 | 4 | 10 | 12 | 24 | 10 | 61 | 28 | 33 | 27 | 31 | 23 | 0 | 32 |
| (TG)6 | 1 | 0 | 1 | 5 | 7 | 11 | 28 | 17 | 38 | 21 | 28 | 31 | 18 | 39 | 0 | 14 |
| (TG)7 | 0 | 1 | 0 | 0 | 2 | 6 | 14 | 4 | 30 | 16 | 14 | 9 | 18 | 21 | 0 | 6 |
| (TG)8 | 1 | 1 | 2 | 1 | 2 | 3 | 10 | 3 | 17 | 5 | 3 | 12 | 13 | 13 | 0 | 4 |
| (TG)9 | 0 | 0 | 0 | 0 | 2 | 3 | 6 | 5 | 8 | 2 | 4 | 7 | 19 | 11 | 0 | 3 |
| (TT)1 | 84 | 81 | 94 | 208 | 315 | 438 | 646 | 448 | 7 | 486 | 671 | 1049 | 874 | 1859 | 0 | 2255 |
| (TT)2 | 2 | 2 | 6 | 14 | 11 | 21 | 32 | 27 | 748 | 22 | 20 | 33 | 41 | 101 | 0 | 100 |
| (TT)3 | 0 | 0 | 0 | 4 | 0 | 7 | 5 | 1 | 38 | 2 | 3 | 4 | 5 | 16 | 0 | 10 |

| | | | | | | | | | | | | | | | | |
|---|---|---|---|---|---|---|---|---|---|---|---|---|---|---|---|---|
| (TT)4 | 0 | 0 | 1 | 0 | 1 | 2 | 2 | 2 | 12 | 0 | 0 | 0 | 0 | 5 | 0 | 2 |
| (TT)5 | 0 | 0 | 0 | 0 | 0 | 0 | 0 | 0 | 3 | 0 | 0 | 0 | 1 | 1 | 0 | 0 |
| (TT)6 | 0 | 0 | 0 | 0 | 0 | 0 | 0 | 0 | 0 | 0 | 0 | 0 | 1 | 0 | 0 | 0 |
